# Tunable differentiation of human CD4^+^ and CD8^+^ T cells from pluripotent stem cells

**DOI:** 10.1101/2024.10.29.620998

**Authors:** Ross D. Jones, Kevin Salim, Laura N. Stankiewicz, John M. Edgar, Lorna Leon, Jana K. Gillies, Ali Murtaza, Lauren J. Durland, Divy Raval, Thristan P. Taberna, Han Hsuan Hsu, Carla Zimmerman, Yale S. Michaels, Fabio M. V. Rossi, Megan K. Levings, Peter W. Zandstra

## Abstract

Allogeneic T cell therapies are a highly desirable option to circumvent the cost and complexity of using autologous T cells to treat diseases. Allogeneic CD8^+^ T cells can be made from pluripotent stem cells (PSCs), but deriving CD4^+^ T cells from PSCs remained a significant challenge. Using feeder-and serum-free conditions, we found that CD4^+^ versus CD8^+^ T cell commitment from PSCs can be controlled by fine-tuning the dynamics of Notch and T cell receptor signaling delivered to CD4^+^CD8^+^ double positive T cells. Notch signaling negatively impacts CD4^+^ T cell commitment, and its timed removal allows generation of clonally-diverse and expandable CD4^+^ T cells from PSCs. The resulting CD4^+^ T cells respond to cytokine-mediated polarization by differentiating into Th1, Th2, or Th17 cells, recapitulating canonical helper cell function. These findings represent a significant step towards using PSC-derived CD4^+^ T cells as a low cost, off-the-shelf, cell therapy.

## Introduction

Engineered T cells have an enormous potential to treat cancer, autoimmunity, and infectious disease^1–3^. However, the high cost and logistical complexity of manufacturing and administering autologous (patient-derived) T cells significantly limit access to T cell therapies. There has therefore been strong interest in scalable, cost-effective production of allogeneic (off-the-shelf) T cells^4–6^. A promising source of allogeneic T cells is human pluripotent stem cells (PSCs), which are capable of indefinite expansion and more amenable to genetic engineering than primary cells, and can be differentiated *in vitro* to cytotoxic (CD8^+^) T cells^7–10^. Recently, we and others identified clinically-relevant, fully-defined (feeder-and serum-free) conditions to differentiate PSCs into CD8^+^ T cells^11–14^. However, a long-standing challenge for PSC-to-T cell manufacturing, especially in feeder-free systems, has been the inability to make substantial numbers of mature helper (CD4^+^) T cells^6^.

CD4^+^ T cells are key orchestrators of the immune system and their absence from PSC-derived T cell populations may limit clinical efficacy. CD4^+^ T cells play an essential role in “helping” CD8^+^ T cells eliminate cancer in the context of both engineered T cell therapies^15–17^ and immunotherapies^18–22^. Moreover, through their ability to differentiate into different helper T (Th) cell subsets, such as Th1, Th2, and Th17 cells, CD4^+^ T cells can orchestrate and regulate diverse immune responses.

Natural differentiation of CD4^+^ vs CD8^+^ T cells is well-studied and primarily driven by the dynamics of T cell receptor (TCR) signaling^23,24^. In the thymus, T cell progenitors develop into double-positive (DP) CD4^+^CD8^+^ cells, then undergo V(D)J recombination and negative and positive selection, ultimately resulting in cells with functional TCRs that complex with CD4 or CD8 co-receptors to recognize peptide-loaded major histocompatibility (MHC) Class I (CD8^+^ T cells) or Class II (CD4^+^ T cells) proteins. Commitment to the CD4 or CD8 lineage is controlled by TCR signaling duration^25,26^; in selecting thymocytes, TCR interactions with MHC Class I generate shorter signals due to transient downregulation of CD8, interrupting TCR-MHC Class I interactions^23,24^. Differences in TCR signaling dynamics lead to selective activation of RUNX3 (CD8) or ThPOK (*ZBTB7B*, CD4), mutually-repressive lineage-defining transcription factors^27–29^.

An outstanding question is why current PSC-to-T cell differentiation protocols lead to poor CD4^+^ T cell development. One explanation could be that differentiating cells have limited MHC Class II expression^30^. However, since most protocols replicate positive selection through MHC-independent TCR stimulation with anti-CD3 antibodies, this seems an unlikely explanation. Rather, we hypothesized that the TCR stimulation regime itself was suboptimal, creating a strong bias towards CD8^+^ T cell development. Of particular interest were effects of Notch signaling, a pathway crucial to T cell development and known to have dose-dependent effects on TCR signaling and CD8^+^ T cell development^31,32^. Notably, in a study using artificial thymic organoids to produce T cells from PSCs^30^, more CD4^+^ T cells developed when using MS5 feeder cells expressing the Notch ligand DLL1 (Delta-like ligand 1) rather than the stronger ligand, DLL4^33^.

We thus studied the combined effects of Notch and TCR signaling levels on *in vitro* differentiation of PSCs to CD4^+^ vs CD8^+^ T cells. We found that optimization of Notch and TCR stimulation enables production of clonally diverse, phenotypically mature, and functional PSC-CD4^+^ T cells, with yields similar to PSC-CD8^+^ T cells. Importantly, these results were obtained using fully-defined media in the absence of feeder cells. The ability to reliably produce controlled proportions of CD4^+^ and CD8^+^ T cells paves the way for improved *in vitro* isogenic immune models and improved off-the-shelf T cell therapies.

## Results

### Notch signaling suppresses production of CD4^+^ T cells from PSCs

*In vitro* production of T cells from PSCs is a multi-stage, ∼6-8 week process that replicates key aspects of *in vivo* T cell development^13,34^ (**Figures 1A-B** and **S1A**). Since TCR signaling strength and duration is a key determinant of CD4^+^ vs CD8^+^ T cell commitment, we first tested the effects of different levels of TCR stimulation on CD4^+^ T cell differentiation from the iPS11 iPSC line (ALSTEM). After generating iPSC-DP cells as previously described^13^, cells were treated with different levels of soluble anti-CD2/3/28 TCR-stimulating complexes and re-plated onto fresh DLL4 (10 μg/mL) + VCAM1 (2.5 μg/mL) (**Figure 1C**). Although we hypothesized that more TCR-stimulating input would increase CD4^+^CD8^−^ (CD4 single-positive, 4SP) cell formation^25^, we were surprised to find the opposite. High levels of anti-CD2/3/28 generated more CD4^−^CD8^+^ (CD8 single-positive, 8SP) cells; a small population of 4SP cells was only observed at lower levels (**Figure 1C**). Among cells expressing CD27, a marker of lineage commitment and maturation^35^ (**Figure S1B**), the ratio of 4SP to 8SP cells was maximized at ∼0.1-0.3% anti-CD2/3/28 (**Figure 1C**, lower); we thus used this optimized range for further protocol development.

**Figure 1:**
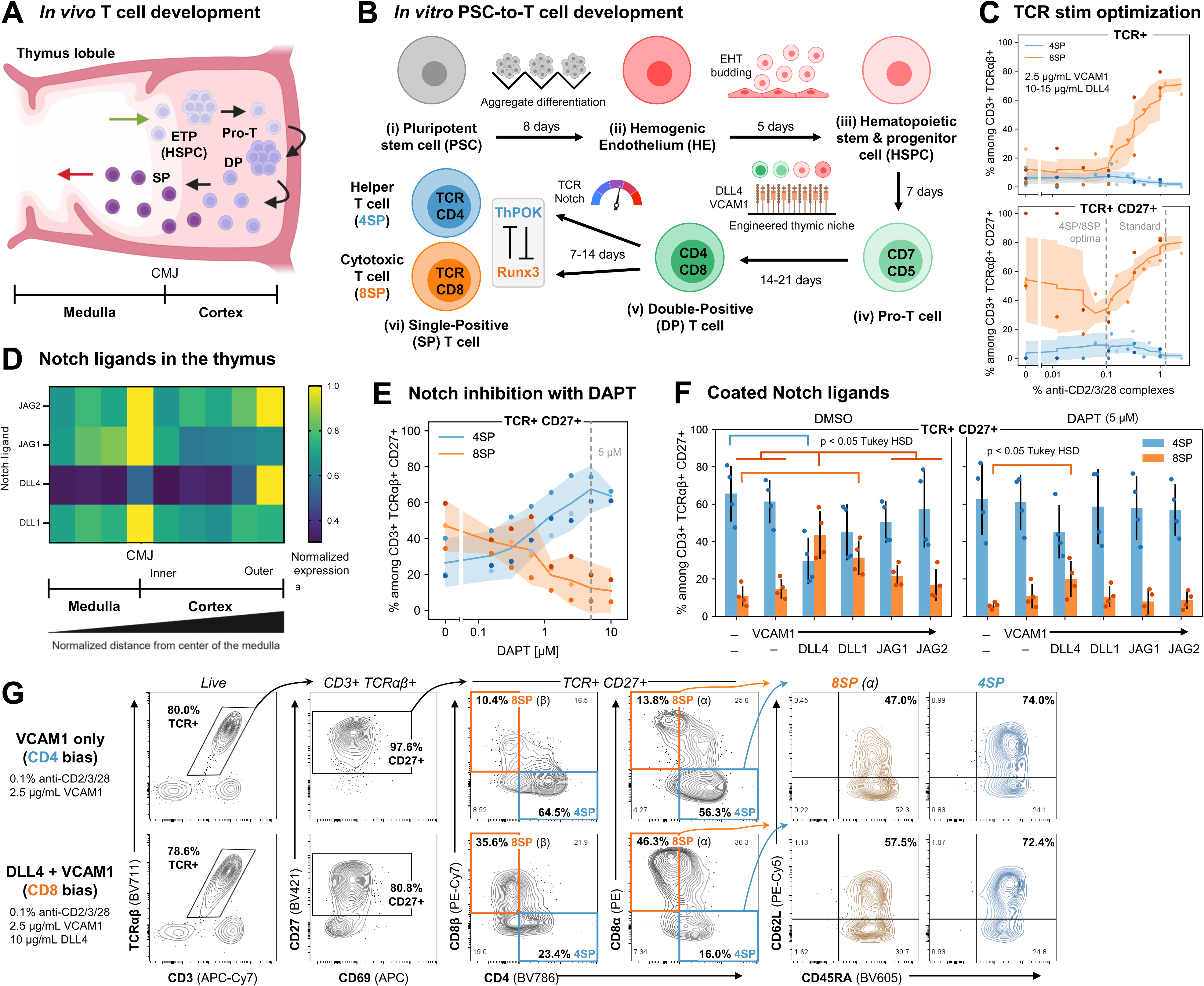
Notch signaling suppresses production of CD4^+^ T cells from PSCs. **(A)** Progression of human T cell development in the thymus. ETP: early thymic progenitor cells; HSPC: hematopoietic stem and progenitor cells; DP: CD4^+^CD8^+^ double-positive cells; SP: CD4 or CD8 single-positive cells. **(B)** Overview of *in vitro* PSC-to-T cell platform. (i) PSCs are aggregated and differentiated to (ii) hemogenic endothelium (HE); (iii) Hematopoietic stem and progenitor cells (HSPCs) bud from HE cells in an endothelial-to-hematopoietic transition (EHT); HSPCs are then sequentially differentiated into (iv) Pro-T, (v) DP, and (vi) SP T cells. This work focuses on the last 1-2 weeks of T cell maturation from DP cells to CD4 or CD8 single positive (4SP, 8SP) cells. ThPOK and RUNX3 are driving transcription factors for CD4^+^ and CD8^+^ T cells, respectively. **(C)** iPSC-derived cells after 2-3 weeks of DP maturation were seeded into wells coated with 10-15 μg/mL DLL4 + 2.5 μg/mL VCAM1 and stimulated with anti-CD2/3/28 to induce positive selection to SP cells, then phenotyped with flow cytometry one week later. Top panel: CD3^+^TCRαβ^+^ cells; bottom panel: more mature CD3^+^TCRαβ^+^CD27^+^ cells (lineage-committed). 4SP: CD4^+^CD8β^−^; 8SP: CD4^−^CD8α^+^; n=4 independent differentiations; dashed lines indicate the standard concentration of anti-CD2/3/28 (1.25%) and optima for increasing the ratio of 4SPs to 8SPs (0.1%). **(D)** Relative expression of different Notch ligands in postnatal thymus (averaged across lines drawn from the center of the medulla to edge of the cortex in n=3 thymus lobules). **(E-F)** iPSC-derived cells after 3 weeks of DP maturation were seeded into wells coated with 10μg/mL Notch ligand +2.5 μg/mL VCAM1 and stimulated with 0.1% anti-CD2/3/28, then measured two weeks later. Notch signaling was inhibited with the γ-secretase inhibitor DAPT (5μM in (F)). n=3 (E) or n=4 (F) independent differentiations. **(G)** Representative examples of cells in (F) differentiated ±DLL4 (both −DAPT), causing CD4 (−DLL4) or CD8 (+DLL4) lineage bias. For panels (C), (E), and (F): media: PSC2; lines + shaded regions: rolling average ± rolling 95% CI; errorbars: standard deviation. See also **Figures S1** and **S2**.

We next examined the effect of Notch signaling on iPSC-DP-to-SP induction. In the neonatal thymus, Notch ligands are at highest densities in the cortical-medullary junction and outer cortex, with diminishing density in the inner cortex, the site of positive selection, especially for the strongest ligand DLL4^36^ (**Figure 1D**). To test if Notch signaling levels affect iPSC-CD4^+^ T cell formation, we stimulated iPSC-DP cells with 0.1% anti-CD2/3/28 in the presence of DLL4 and VCAM1, then inhibited Notch signaling using the gamma secretase inhibitor (GSI) DAPT. With increasing DAPT, we observed a striking dose-dependent switch from 8SP to 4SP cell induction (**Figure 1E**), suggesting that Notch signaling biases *in vitro* PSC-to-T cell differentiations towards CD8^+^ T cells.

Different Notch ligands have different strengths of Notch pathway activation^33,37,38^, so we next tested whether the CD8-biasing effect of Notch was DLL4-specific. iPSC-DP cells were transferred to plates coated with different Notch ligands (DLL4, DLL1, JAG1, JAG2, or none), and stimulated with 0.1% anti-CD2/3/28. We found increasing CD8 lineage bias for DLL4 > DLL1 > JAG1 ≈ JAG2, correlating with the known affinity and activation potential of each Notch ligand^33^ (**Figure 1F**, left). Inhibiting Notch pathway activation with DAPT restored CD4 lineage bias for cells cultured on all Notch ligands (**Figure 1F**, right).

Removing Notch signaling resulted in a large population of CD4^+^ T cells that were negative for both CD8α and CD8β, and expressed several markers of mature thymocytes and naïve T cells including CD27, CD45RA, and CD62L (**Figure 1G**). Maturation was similar for cells grown on different Notch ligands, but with a lower rate of CD27 induction for DLL4 (**Figure S1C**). Interestingly, we observed that CD8β was downregulated faster than CD8α in iPSC-T cells that committed to the CD4 lineage (**Figure S1D**). Comparing TCR^+^CD27^+^ cells 7 vs 14 days after stimulation, the number of CD4^+^CD8β^−^ 4SP cells at day 7 predicted the numbers of CD4^+^CD8β^−^ and CD8α^−^ 4SP cells one week later (R^2^ = 0.74 and 0.76, respectively). Thus, early committed iPSC-CD4^+^ T cells can be identified as CD3^+^TCRαβ^+^CD27^+^CD4^+^CD8β^−^ 4SPs even before CD8α downregulation.

We next tested if the PSC-CD4^+^ T cell induction conditions are generalizable to different media conditions and cell sources. First, we applied our optimized Notch and TCR stimulation regimes to iPSC-DP cells grown in different compositions of serum-free media (**Figures S2A-B**). In all cases, we confirmed robust induction of the CD4 lineage (**Figure S2C**). We then differentiated the H1 human embryonic stem cell line to Pro-T and DP cells and confirmed subsequent induction to CD4^+^ T cells using our optimized conditions (**Figure S2D**).

### Tuning Notch and TCR stimulation controls the ratio of iPSC-CD4^+^ vs CD8^+^ T cells

The proportion of CD4 to CD8 T cells is an important factor in the efficacy of CAR-T cell therapy^16,39^, so we next tested whether we could control the ratio of iPSC-CD4^+^ vs CD8^+^ T cells in a single culture by modulating levels of both Notch and TCR stimulation during the DP-to-SP transition (**Figure 2A**). To do so, we seeded cells onto wells coated with different levels of DLL4 (0-10 μg/mL, all with 2.5 μg/mL VCAM1) and treated with different levels of TCR stimulating reagents. We hypothesized that different modes of T cell activation might influence CD4 vs CD8 lineage bias. We therefore tested the effects of stimulating iPSC-DP cells with phytohemagglutinin (PHA), phorbol myristate acetate (PMA) and ionomycin (Iono), or anti-CD2/3/28 as above (**Figure 2A**). PHA is a lectin that generally cross-links glycosylated cell surface proteins, including the TCR; whereas PMA is an analogue of diacyl glycerol (a TCR signaling intermediate molecule), often used with Iono, a calcium ionophore, to recapitulate TCR signaling in a TCR-independent manner.

**Figure 2:**
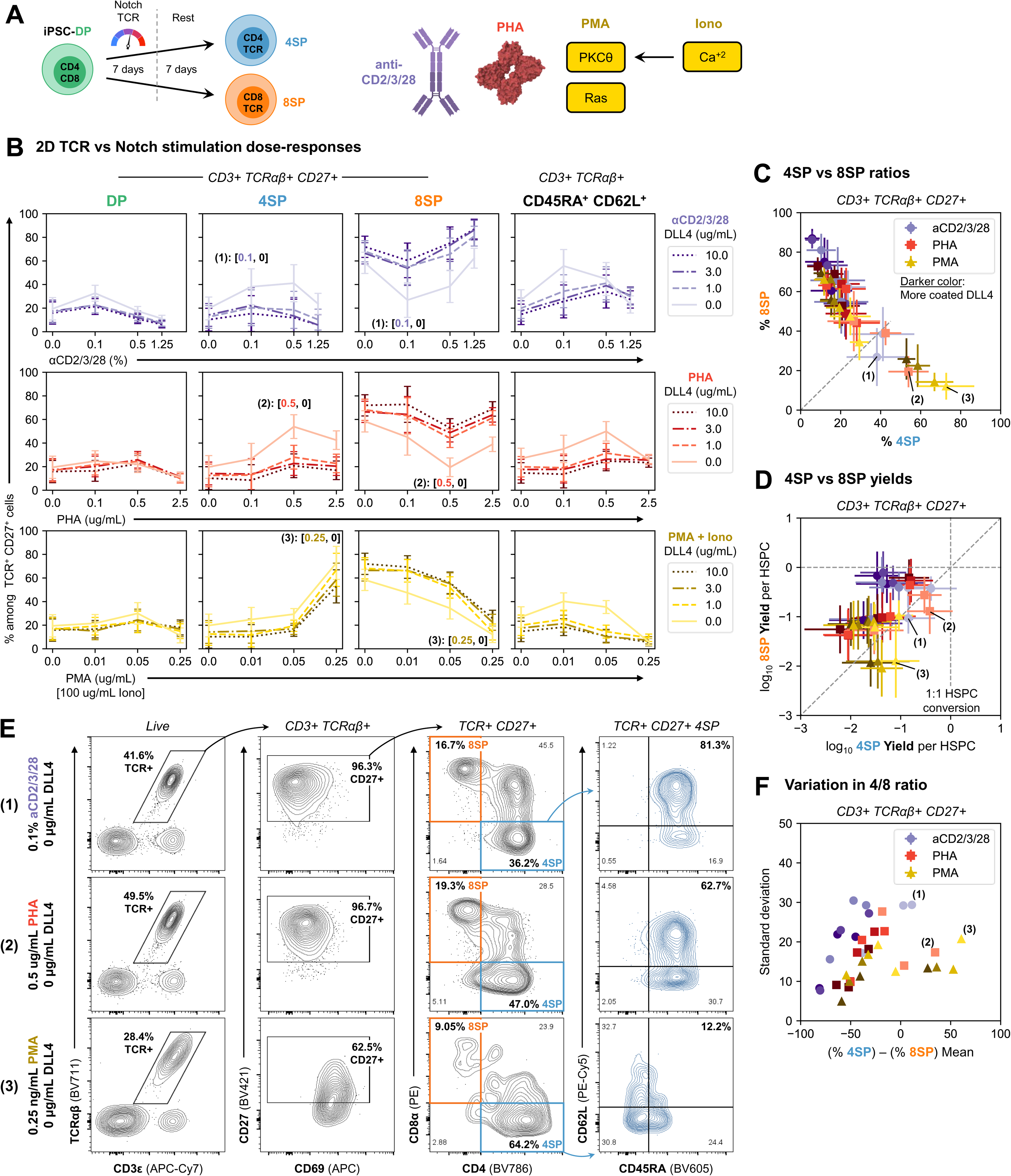
Tuning Notch and TCR stimulation controls the ratio of iPSC-CD4^+^ vs CD8^+^ T cells. **(A)** iPSC-derived cells after 3 weeks of DP maturation were seeded into wells freshly coated with different levels of DLL4 + 2.5 μg/mL VCAM1 and stimulated with different levels of TCR pathway activators to induce positive selection to SP cells. Cells were fed three days later without additional TCR stimulation, then rested for a week by passaging onto wells coated with VCAM1 only and in media without TCR stimulation. Anti-CD2/3/28 and PHA both crosslink the TCR with other receptors to induce multiple downstream signaling pathways; Phorbol-myristate-acetate (PMA) is a DAG agonist and binds to and activates PKCθ and Ras. Ionomycin (Iono) is an ionophore that binds Ca^+2^ and increases its cytoplasmic concentration. PMA+Iono together directly actuate downstream signaling pathways independent of the TCR receptor. **(B)** Percent of CD3^+^TCRαβ^+^CD27^+^ cells that stayed DP or converted to 4PS vs 8SP. The rightmost column shows the percent of TCR^+^ cells that matured to a CD45RA^+^CD62L^+^ naïve-like T cell state. Each line represents a different level of coated DLL4. The labels (1), (2), and (3) highlight conditions that optimize the ratio of 4SP to 8SP cells across panels (B-F). 4SP: CD4^+^CD8β^−^; 8SP: CD4^−^CD8α^+^; media: PSC2; n=4 independent differentiations; errorbars: mean ± standard deviation. **(C, D)** Comparison of the percent (C) and yield per CD34^+^CD43^+^ HSPC input (D) of TCR^+^CD27^+^ 4SP vs 8SP cells across all conditions with non-zero TCR stimulation input levels. Darker colors indicate increasing concentrations of coated DLL4. **(E)** Representative flow plots for optimal 4SP-inducing conditions (1), (2), and (3) for different TCR stimulating reagents. **(F)** Comparison of the mean and standard deviation in the percent of cells that commit to CD4^+^ vs CD8^+^ T cells across all conditions. See also **Figure S3**.

Similar to the response with anti-CD2/3/28, there was an optimal PHA or PMA+Iono dose for induction of 4SP cells. In all cases, decreasing DLL4 increased the percent and yields of 4SP cells (**Figures 2B** and **S3A**). In the absence of stimulating inputs, there was a strong bias in spontaneous differentiation towards 8SP cells. Notably, the proportion of TCRαβ^+^ cells that acquired a mature phenotype (CD45RA^+^CD62L^+^) also increased with decreasing DLL4 (**Figures 2B** and **S3B**), corresponding with our prior observation that DLL4 suppressed expression of the maturation marker CD27 (**Figure S1C**). For PHA, the overall cell phenotype pattern was similar to those stimulated with anti-CD2/3/28, with both reagents generating mature CD4^+^ T cells. Interestingly, the optimal dose of PMA for CD4^+^ T cell induction (0.25 ng/mL) gave the lowest percent of CD45RA^+^CD62L^+^ cells, suggesting reduced maturation.

Overall, varying cell activation conditions enabled production of a broad range of 4SP:8SP ratios (**Figure 2C**). PMA+Iono stimulation resulted in the highest proportion of 4SPs, but with lower yields (**Figures 2D-E** and **S3A**). Notably, there were many conditions with high yields of both 4SPs and 8SPs (**Figure 2D**). Conditions that generated intermediate 4SP:8SP ratios showed the highest variance in the proportion of 4SP vs 8SP cells in different experimental replicates (**Figure 2F**). This variation was highest with anti-CD2/3/28 and lowest with PMA, possibly indicating sensitivity to factors such as cell density under certain stimulation conditions.

Collectively, these results indicate that tunable and predicable ratios of CD4^+^ vs CD8^+^ T cells can be generated by controlling Notch and TCR pathway stimulation. Importantly, optimal levels of multiple TCR pathway stimulators are compatible with PSC-CD4^+^ T cell production. Overall, these results suggest removal of Notch signaling and optimization of TCR stimulation is a broadly-applicable strategy to enhance CD4^+^ T cell production from PSCs.

### iPSC-CD4^+^ T cells are mature and expandable *in vitro*

To further evaluate our PSC-CD4^+^ T cell induction protocol, we investigated expression of naïve T cell markers and the lineage-defining transcription factors ThPOK (CD4) and RUNX3 (CD8). iPSC-DP cells were cultured in conditions optimized to remain DP or differentiate to CD4^+^ or CD8^+^ T cells using either anti-CD2/3/28 or PHA (**Figure 3A**) and one week later the expected cell type bias was confirmed (**Figure 3B**). Compared to DP cells, newly committed 4SPs and 8SPs upregulated CD27, CD45RA, CD45RO, and CCR7 (**Figure 3C**). Surprisingly, CD62L levels were already high in DPs and decreased in both 4SPs and 8SPs (**Figure 3C**). As expected, 4SPs upregulated ThPOK but not RUNX3, with the opposite pattern seen for 8SPs (**Figure 3C**). The pattern of naïve cell markers and lineage transcription factors was not significantly different between cells stimulated with anti-CD2/3/28 or PHA.

**Figure 3:**
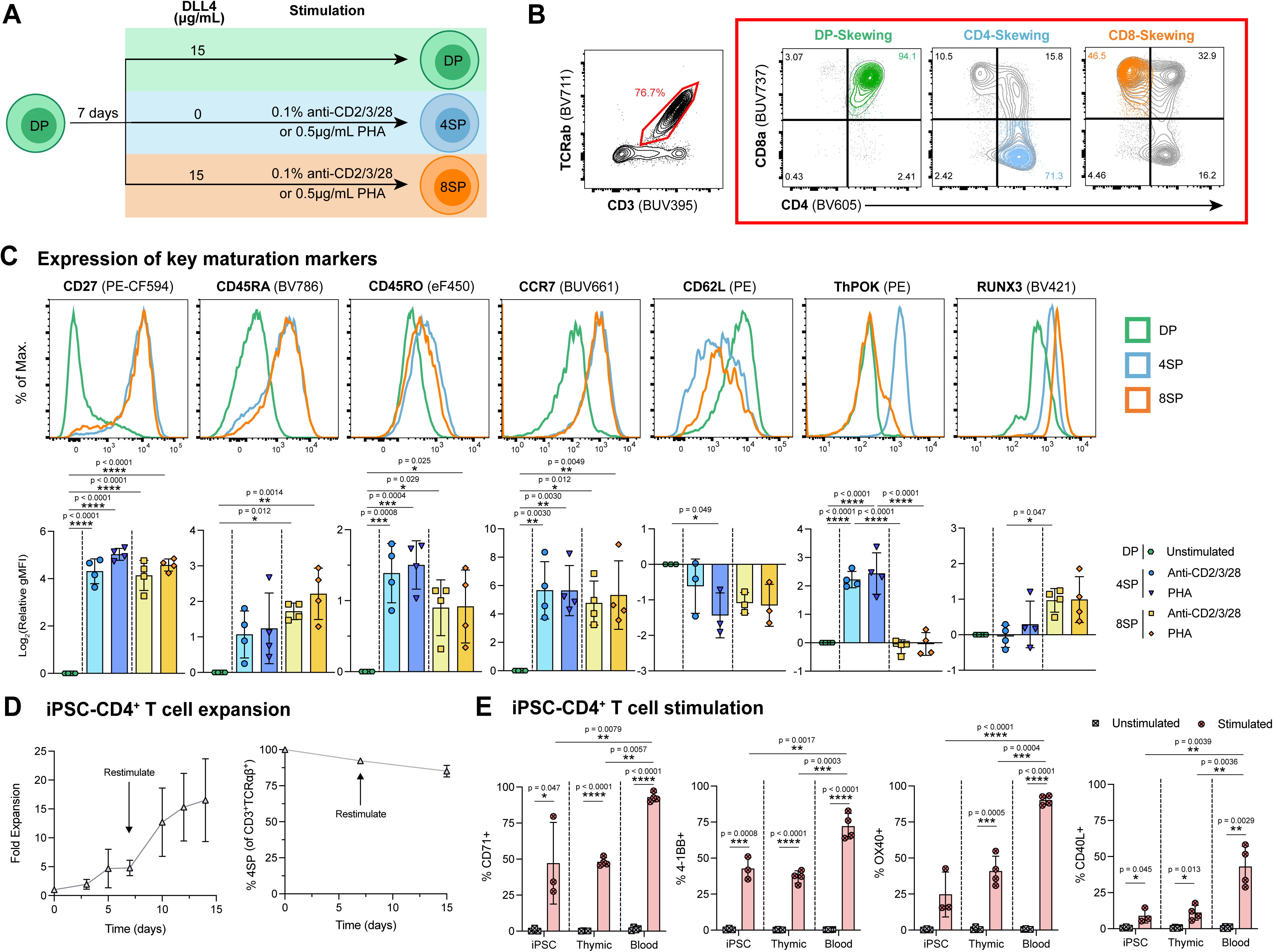
iPSC-CD4^+^ T cells are mature and expandable *in vitro*. **(A)** iPSC-DPs were transferred into different downstream conditions in StemSpan Maturation media: DP-skewing: 15 mg/mL DLL4 and no stimulation; CD4-skewing: 0 mg/mL DLL4 and stimulation with 0.1% anti-CD2/3/28 or 0.5 μg/mL PHA-L; or CD8-skewing: 15 mg/mL DLL4 and stimulation with 0.1% anti-CD2/3/28 or 0.5 μg/mL PHA-L. Expression of naïve CD4^+^ T cell markers and lineage-specific transcription factors were measured after 7 days by flow cytometry. **(B)** Representative flow plots of CD3^+^TCRab^+^ cells and their CD4 vs CD8a expression pattern after stimulating iPSC-DPs for 7 days. **(C)** Top: Representative histograms of naïve CD4^+^ T cell markers and lineage-specific transcription factors of iPSC-DPs, -4SPs, and -8SPs after 7 days of stimulation; Bottom: log_2_-transform of each marker’s relative geometric mean fluorescence intensity (gMFI) of iPSC-4SP/-8SP normalized to iPSC-DPs. Data is shown with mean ± standard deviation; n=3-4 independent differentiations; statistical significance was determined using ordinary one-way ANOVA with p values shown. **(D)** Sorted iPSC-CD4^+^ T cells were expanded for 14 days in Immunocult-XF + 100 IU/mL IL-2, with stimulation on day 0 and 7 using 2.5% anti-CD2/3/28. The left plot shows the fold-expansion and the right plot shows the percent 4SP cells (CD4^+^CD8α^+^ within CD3^+^TCRab^+^) throughout expansion. Data is shown with mean ± standard deviation; n=4 independent differentiations. **(E)** Expanded cells were rested overnight in 10 IU/mL IL-2, restimulated with anti-CD2/3/28 for 48 hours, and measured for T cell activation markers using flow cytometry. Data is shown with mean ± standard deviation; n=3-4 independent differentiations, or n=3-4 thymic-or blood-donors collected in 2-3 experiments. Statistical significance was determined using unpaired t-tests and ordinary one-way ANOVA with p values shown. See also **Figure S4**.

Next, we investigated if iPSC-CD4^+^ T cells can be expanded while retaining their 4SP phenotype. iPSC-DPs were induced to 4SPs with PHA stimulation for 7 days, then PHA was removed and cells were transitioned into media commonly used to expand primary T cells (ImmunoCult-XF T cell expansion media supplemented with 100 IU/mL IL-2). After 7 days, 4SPs were sorted (**Figure S4A**) and stimulated with anti-CD2/3/28 in the same media. Over the course of 14 days, with restimulation at day 7, the cells proliferated ∼15-fold while retaining the 4SP phenotype in >80% of cells (**Figures 3D and S4B**).

Finally, we assessed if iPSC-CD4^+^ T cells upregulated relevant markers of activation and co-stimulatory function after 48h of stimulation with anti-CD2/3/28. As a comparator, CD4^+^ T cells isolated from postnatal thymus or adult peripheral blood were tested in parallel (**Figures S4C-D**). We found that iPSC-CD4^+^ T cells significantly upregulated all T cell activation markers tested (CD71, 4-1BB, OX40, and CD40L) (**Figures 3E and S4E**). Interestingly, the proportion of marker expression was more similar to that of thymic-CD4^+^ T cells than adult blood-CD4^+^ T cells. Overall, similar to naturally-developed T cells, iPSC-CD4^+^ T cells expanded *in vitro*, retained their CD4 phenotype, and upregulated relevant markers of activation and co-stimulation.

### Notch suppresses TCR signaling to bias towards CD8^+^ T cells

To better understand the mechanism by which Notch and TCR input signals control CD4 vs CD8 lineage commitment in PSC-T cells, we measured key transcription factors GATA-3, ThPOK, and RUNX3 during two weeks of *in vitro* PSC-DP-to-SP induction (**Figure 4A**). iPSC-derived cells three weeks into maturation (day M21) were stimulated with six different combinations of two TCR stimulation levels and three Notch inputs, each with different lineage biases: either low (0.3%, CD4-biased) or high (1.25%, CD8-baised) anti-CD2/3/28; −DLL4/−DAPT (CD4-biased), +DLL4/−DAPT (CD8-biased) or +DLL4/+DAPT (CD4-biased) (**Figure 4B** and **Figure S5A**).

**Figure 4:**
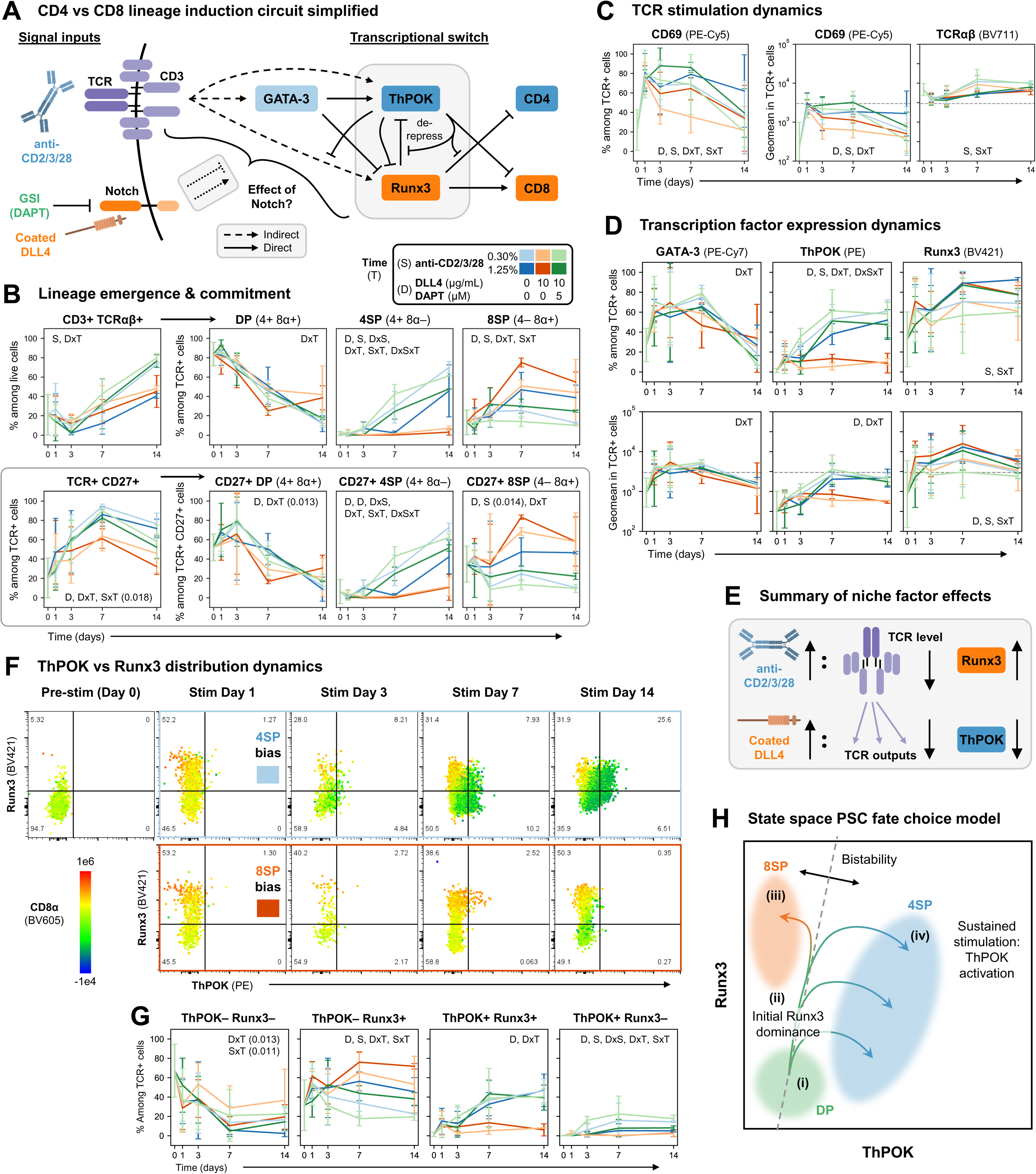
Notch suppresses TCR signaling to bias towards CD8^+^ T cells. **(A)** Simplified CD4 vs CD8 lineage induction circuit^27,46,68–70^. Solid vs dashed lines indicate direct vs indirect interactions, respectively. This experiment aimed to uncover the effect of Notch on the key transcription factors GATA-3, ThPOK, and Runx3 PSC-T cell maturation. **(B-D)** Dynamics of (B) TCR, CD27, CD4, and CD8 expression, (C) TCR signaling, and (D) transcription factors in different TCR and Notch stimulation conditions. Dashed lines in geomean plots indicate positive gate threshold. **(E)** Summary of how anti-CD2/3/28 and Notch inputs affect downstream gene expression to bias CD4 vs CD8 lineage commitment. **(F-G)** RUNX3 vs ThPOK expression overlayed with CD8α expression in key CD4 vs CD8 biased conditions (F) and compared across conditions (G). **(H)** Model of cell progression through ThPOK vs RUNX3 state space during iPSC-DP-to-SP induction. When induced, ThPOK dominates a bistable switch with Ruxn3 by repressing RUNX3 (indirectly), activating itself (indirectly) and preventing RUNX3 from repressing itself (directly). Cells at the boundary between ThPOK and RUNX3 are bistable and can commit to either lineage. For panels (B), (C), (D), and (G): errorbars: mean ± standard deviation; n=4 independent differentiations (n=2 at day S14); media: StemSpan Mat + JAC Ultra; Inset indicators D, S, and T refer to significance of DLL4/DAPT variants (D), anti-CD2/3/28 levels (S), and interactions (x) with each other and time (T) using three-way ANOVA with a significance threshold of p < 0.01; p-values close to but above the significance threshold are shown. Detailed statistics are provided in **Table S1**. See also **Figure S5**.

Following stimulation (all conditions), TCR expression sharply decreased over the first 1-3 days, then rebounded over the rest of the timecourse (**Figure 4B**, top left). Among iPSC-derived CD3^+^TCRαβ^+^ cells, CD4^+^CD8α^+^ DPs gradually converted to different ratios of CD4^+^CD8α^−^ 4SPs or CD4^−^CD8α^+^ 8SPs depending on the condition (**Figure 4B**, top right). In conditions +DLL4/−DAPT, 4SP induction was suppressed, nearly half the cells remained DP, and TCR reactivation was reduced. From the kinetic signaling model, we expected to see rapid downregulation of CD8^24^, but this was only observed for CD8β: CD4^+^CD8β^−^ 4SPs appeared within one day in conditions −DLL4 or +DAPT (**Figure S5B**).

Consistent with our prior observations (**Figures S1C**, **2B**, and **S3C**), CD27 induction was reduced by ∼20-30 percentage points in conditions +DLL4/−DAPT 7-14 days post-stimulation (**Figure 4B**, bottom left). The dynamics of CD27^+^ DP/4SP/8SP populations mirrored their broader TCR^+^ subsets (**Figures 4B** and **S5B**, bottom right). Strikingly, committed CD27^+^CD4^+^CD8β^−^ 4SPs appeared within one day of stimulation and were remarkably stable throughout the timecourse for conditions −DLL4/+DAPT (**Figure S5B**, bottom right). This finding is consistent with (i) our prior observation of PSC-derived CD27^+^CD4^+^CD8β^−^ 4SPs appearing earlier than CD27^+^CD4^+^CD8α^−^ 4SPs (**Figure S1D**), (ii) the notion that CD27 marks stably committed cells^35^, and (iii) recent work indicating CD4 lineage commitment is much more rapid than CD8 lineage commitment^40^.

Like CD27, the TCR stimulation marker CD69 was also reduced in +DLL4/−DAPT conditions (**Figure 4C**). Conditions with high anti-CD2/3/28 input (1.25%) induced higher CD69 expression but slowed the post-stimulation recovery of TCR expression (**Figure 4C**). This may be due to stronger stimulation input triggering more negative feedback via TCR ubiquitination and downregulation^41,42^.

Given the large differences in CD69 activation in +DLL4/−DAPT conditions, we were surprised to see only minor differences in the dynamics of GATA-3, a pro-CD4 lineage transcription factor that is induced by TCR stimulation^43^, activates ThPOK^44^, and represses RUNX3^45^. In all conditions, GATA-3 expression rapidly increased within one day of stimulation, then slowly decreased between days 3-14 (**Figure 4D**, left). The decrease was faster in +DLL4/−DAPT conditions, potentially due to reduced TCR pathway activation. Conversely, ThPOK expression was almost completely blocked in Notch-stimulating conditions (**Figure 4D**, middle), directly linking to the CD8 lineage bias of Notch in our system. Unexpectedly, we observed rapid and strong induction of RUNX3 in all conditions, with higher expression in conditions with higher (1.25%) anti-CD2/3/28 input (**Figure 4D**, right). This contrasts with *in vivo* DP-to-SP transitions, where RUNX3 activation is delayed^27^. Early expression of RUNX3 may explain the lack of CD8α downregulation following stimulation.

Overall, our results in **Figure 4B-D** show that high levels of anti-CD2/3/28 input drive down TCR expression and increase RUNX3 expression, contributing to CD8 lineage bias (**Figure 4E**). Strong Notch signaling by DLL4 reduces downstream TCR signaling pathway outputs, especially ThPOK, thereby suppressing the CD4 lineage (**Figure 4E**).

Looking at the joint expression distribution of ThPOK and RUNX3, we were surprised to see that most ThPOK^+^ cells also expressed RUNX3 (**Figures 4F-G**). ThPOK expression correlated directly with CD4 expression, whereas many RUNX3^+^ cells were CD8α^−^ (**Figures S5C-D**). RUNX3^+^CD8α^−^ cells expressed high levels of ThPOK (**Figure S5D**), consistent with ThPOK’s ability to self-activate, prevent RUNX3 silencing of the *ThPOK* and *CD4* genes, and suppress the *CD8* locus^46^. The gradual increase in ThPOK levels over time (**Figures 4D,F**) compared to the more rapid commitment to the CD4 lineage inferred by CD4 vs CD8β expression in TCR^+^CD27^+^ cells (**Figure S5B**) is consistent with ThPOK’s reported role in “sealing” rather than initiating CD4 fate^47^. The expression level of CD8α corresponded well with the ratio of ThPOK vs RUNX3 in the cell population, consistent with ThPOK and RUNX3 proteins competing to repress and activate the *CD8* locus, respectively^48,49^.

To explain our observations, we put forward a state space model of PSC-derived T cell commitment (**Figure 4G**). Starting as PSC-DPs (i), stimulated cells initially up-regulate RUNX3 (ii). Differences in TCR and Notch input levels translate to RUNX3 (iii) or ThPOK dominance (iv), leading to CD8 vs CD4 lineage bias, respectively (**Figure 4G**). At a critical boundary ratio of ThPOK vs RUNX3 levels, the system is bistable and cells can stochastically commit to either CD4 or CD8 lineage; leading to higher variability in the percent of 4SP vs 8SP cells induced in equal bias conditions (**Figure 2F**).

### iPSC-CD4^+^ T cells express helper T cell genes

To further characterize iPSC-CD4^+^ T cells, we performed CITE-seq and scTCR-seq on iPSC-T cells from a differentiation that produced an approximately equal ratio of 4SP and 8SP cells (**Figure 5A**). Single cell clustering confirmed that the cultures contained CD4^+^ and CD8^+^ T cells, along with quiescent and proliferating DPs and several populations of innate-like T cells, including ILC3-like cells and γδ T cells (**Figure 5B**). Focusing on αβ T cell populations, cells in the CD4_mature cluster expressed maturation markers seen by flow cytometry (**Figure 3**), such as *CD27*, *CCR7*, and *CD40LG*, as well as others including *CCR4*, *IL7RA*, *S1PR1*, and *KLF2*. Cells in the CD4_mature cluster also expressed CD4 lineage induction-associated genes including *LEF1* and *TCF7* (TCF-1) and lacked expression of CD8^+^ T cell-specific genes such as *RUNX3*, *CXCR6*, *GZMB*, *GZMK*, and *PRF1* (**Figures 5C** and **S6A-B**). The DP clusters contained cells expressing a mix of CD4-and CD8-associated genes, likely due to low cluster separation. We confirmed with a separate scRNAseq + CITE-seq dataset containing pre-stimulation DP cells that unsignaled DPs lacked expression of maturation and lineage-specific markers (**Figure S6C-D**).

**Figure 5:**
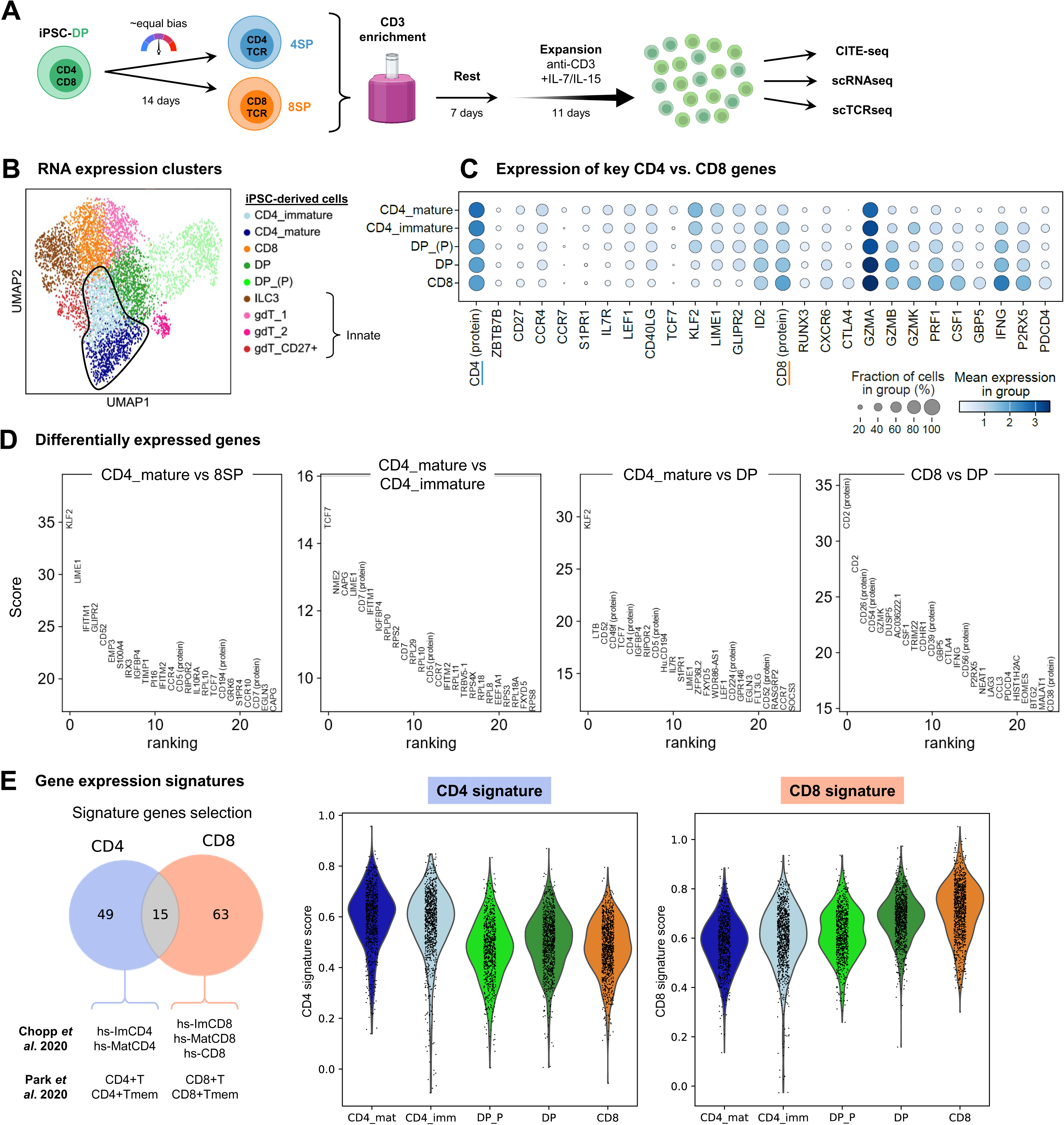
iPSC-CD4^+^ T cells express helper T cell genes. **(A)** iPSC-derived cells at two weeks of DP maturation were induced to SP cells over two weeks in conditions with approximately equal bias (0.1% [week 1], then 0.3% [week 2] anti-CD2/3/28 on 10 μg/mL DLL4 + 2.5 μg/mL VCAM1 [both weeks]). CD3^+^ cells were then magnetically enriched, rested for one week, and expanded on CD3 [clone OKT3] + RetroNectin in IL-7/IL-15-based media^13,56^. After 11 days of expansion, the cells were collected and measured with CITE-seq, scRNAseq, and scTCRseq. **(B)** UMAPs and RNA leiden-based clusters identified immature/mature CD4^+^ T cells, CD8^+^ T cells, DP cells (P: proliferating), ILC3-like cells, and γδ T cells. **(C)** Expression of key maturation, lineage-specific, and differentially-expressed genes in the CD4, CD8, and DP populations. **(D)** Top differentially expressed genes among key cell type clusters. **(E)** CD4 and CD8 signature scores derived from human scRNAseq CD4-or CD8-annotated T cell clusters from Chopp *et al*.^47^ and Park *et* al.^50^. Genes present in both the collated CD4 and CD8 sets were dropped to get the final signatures, which are provided in **Table S2**. See also **Figure S6**.

Comparing differentially-expressed genes (DEGs) between the CD4_mature and CD8 clusters, cells in the former expressed higher levels of signaling proteins such as *LIME1, IFITM1, GLIPR2,* and *CD52*, while cells in the CD8 cluster expressed more inflammatory genes such as *CSF1, GBP5* (IFNγ-induced), IFNγ (*IFNG*), and *P2RX5,* and inhibitory genes such as *CTLA4* and *PDCD4* (**Figure 5C-D**). Comparing DEGs between CD4_mature and CD4_immature clusters, cells in the former expressed more *TCF7*, *NME2*, *CAPG*, and *LIME1*, reflecting changes in metabolism, cell mobility, and TCR signaling as the cells mature.

Finally, we systematically compared the iPSC-derived cells to CD4^+^ and CD8^+^ T cell gene signatures generated from human thymocyte scRNAseq data^47,50^. As expected, cells in the CD4_mature and immature clusters scored highest on the CD4 signature while cells in the CD8 cluster scored highest on the CD8 signature (**Figure 5E**). Overall, iPSC-CD4^+^ and -CD8^+^ T cells take on unique transcriptional profiles that correspond with expected human thymocyte reference transcriptomes.

### iPSC-CD4^+^ T cells express a diverse TCR repertoire

The ability to produce CD4^+^ T cells from PSCs enabled us to perform a comparison of TCR repertoires in iPSC-derived CD4^+^ vs CD8^+^ T cells. For *in vivo* references, we used human thymocyte^50^ and peripheral blood mononuclear cell (PBMC)-derived T cells^51^. First, we confirmed that cells in the iPSC-DP, -CD4, and -CD8 clusters express diverse TCR sequences, though less diverse than thymocytes and PBMC T cells (**Figure 6A**). This may be due to the cell expansion step prior to sequencing (**Figure 5A**), which can promote oligoclonality. As with recent studies of PSC-CD8^+^ T cells by our lab and others^13,14,30^, all iPSC-derived cells expressed TCRs with short fetal thymocytes-like CDR3 lengths (**Figure 6B**), indicating low TdT levels during TCR rearrangement and positive selection^52^. Smaller CDR3 lengths likely contribute to lower diversity in iPSC-derived cells, as in fetal thymocytes.

**Figure 6:**
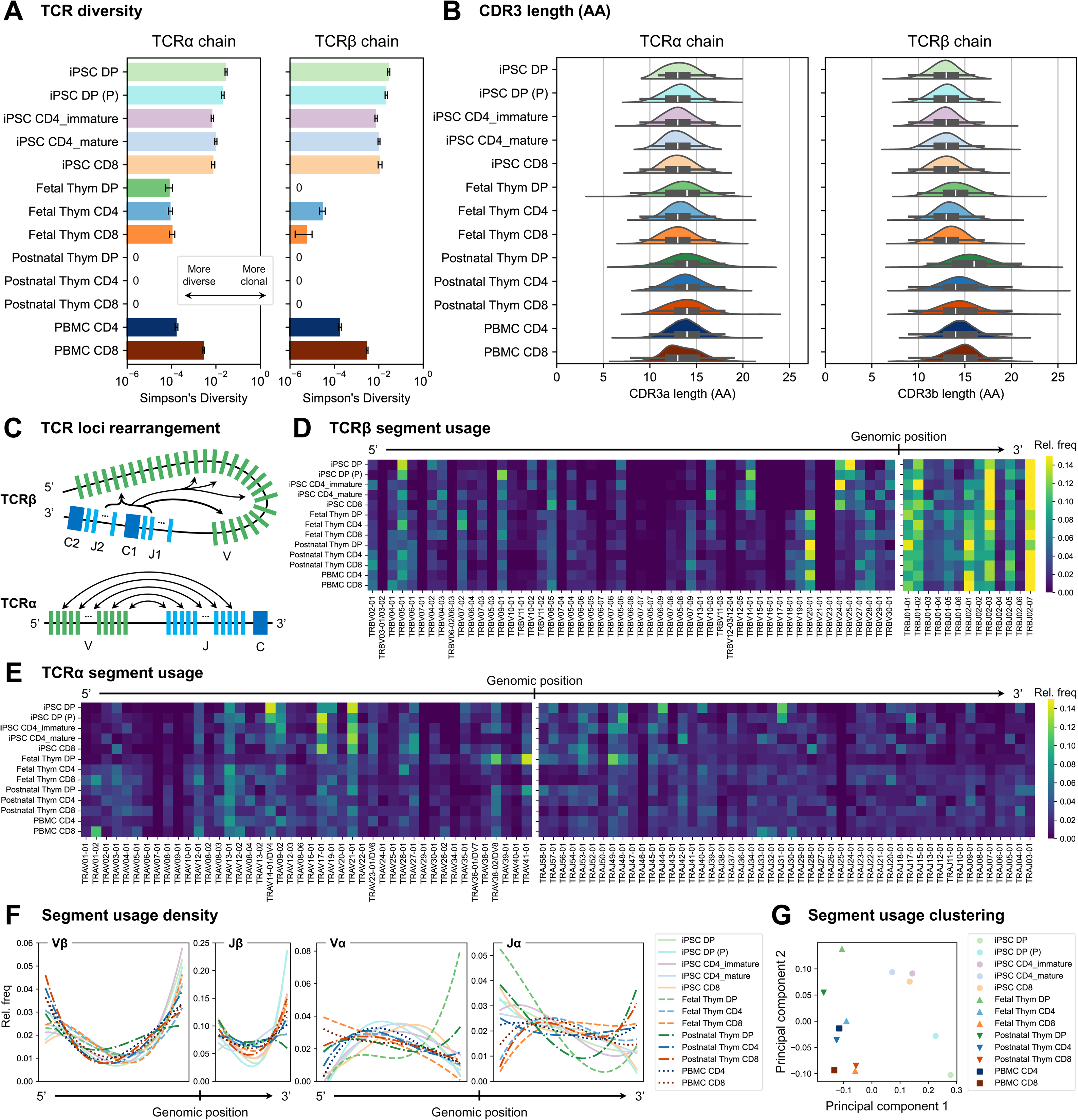
iPSC-CD4^+^ T cells express a diverse TCR repertoire. **(A)** Comparison of TCR diversity, measured by –Simpson’s Diversity, computed on TCR clone counts (unique CDR3 + V + J rearrangements). Values are in the range [0, 1], with lower numbers indicating higher diversity and higher numbers indicating higher clonality. Errorbars: unbiased estimation of standard deviation^71^. **(B)** Comparison of CDR3 sequence length (in amino acids). **(C)** TCR loci rearrangement is positionally-biased (adapted from Park *et al*.^50^); TCR β chain rearrangements initially prefer V segments at the ends of the Vβ array in early developmental stages (Pro-T), and access interior V segments in later stages (DP, SP); α chain rearrangements initially prefer 3’ Vα and 5’ Jα segments in earlier stages (Pro-T, DP), and access increasingly dispersed segments through serial recombination during SP induction and positive selection. **(D, E)** TCRβ (D) and TCRα (E) chain segment usages across iPSC-, thymic-, and blood-T cells. Segments are ordered based on genomic position, frequencies of each row sum to 1. **(F)** B-spline smoothed segment usage densities for each array.

We next analyzed TCR V(D)J rearrangement patterns, which show stage-specific biases in primary thymocytes and are hypothesized to be driven by genomic folding and proximity^50,53^ (**Figure 6C**). The TCR repertoire of iPSC-CD4^+^ and CD8^+^ T cells both used diverse V and J segments across the entire TCRα and β loci (**Figures 6D-E**). Notably, all iPSC-T cells showed a unique pattern of Vβ chain usages for several segments near the center of the Vβ array, in the range TRBV07-08 to TRBV25-01 (**Figure 6D**), but the overall density of Vβ chains was otherwise similarly dispersed throughout the array (**Figure 6F**). All iPSC-T cells except the CD4_mature cluster infrequently used middle Jβ segments compared to primary cell types (**Figures 6D** and **6F**).

TCRα chain selection was similar between iPSC-T cells and all primary cell types except fetal DP thymocytes (**Figure 6E**). iPSC-T cell Vα patterns were highly enriched for fragments near the center of the Vα array and were most similar to patterns in postnatal thymocyte-and PBMC-CD4^+^ T cells (**Figures 6E** and **6F**). iPSC-T cell Jα patterns were enriched for fragments near the 5’ end of the Jα array and were most similar to patterns in postnatal thymocyte-DPs and -CD4^+^ T cells (**Figures 6E** and **6F**).

Finally, we clustered each cell type using principal component analysis (PCA) on the relative usage frequency of all V and J segments for both α and β chains (**Figure 6G**). Primary CD4^+^ and CD8^+^ T cells clustered separately, as expected^50^, while iPSC-CD4^+^ and CD8^+^ T cells clustered together and apart from the primary cell types. Overall, these data point to a unique TCR rearrangement process in iPSC-T cells that is similar between iPSC-CD4^+^ and CD8^+^ T cells.

### iPSC-CD4^+^ T cells can polarize into functionally distinct T helper subsets

A hallmark of CD4^+^ T cells is their ability to respond to a specific cytokine milieu and be polarized into specialized, helper subsets which direct and enhance the function of other immune cells. To test if iPSC-CD4^+^ T cells could acquire helper cell functions, we stimulated them with anti-CD2/3/28 in the presence of Th1-, Th2-, or Th17-polarizing cytokines for 7-14 days. As primary cell references, we polarized naïve CD4^+^ T cells from postnatal thymus or PBMCs in parallel. After polarization, cells were tested for acquisition for Th cell lineage-associated phenotypes, including transcription factors, chemokine receptors and cytokines (**Figure 7A**). Under Th0, Th1, and Th2 conditions, expression of CD4 and CD8a was stable across cell sources (iPSC, thymic, and blood), with a small population of DPs present in both iPSC-and thymic-CD4^+^ T cells. However, under Th17 conditions, a significant fraction of DN and CD8^+^ T cells emerged in iPSC-T cells, but not thymic or blood cells (**Figure S7A-C**).

**Figure 7:**
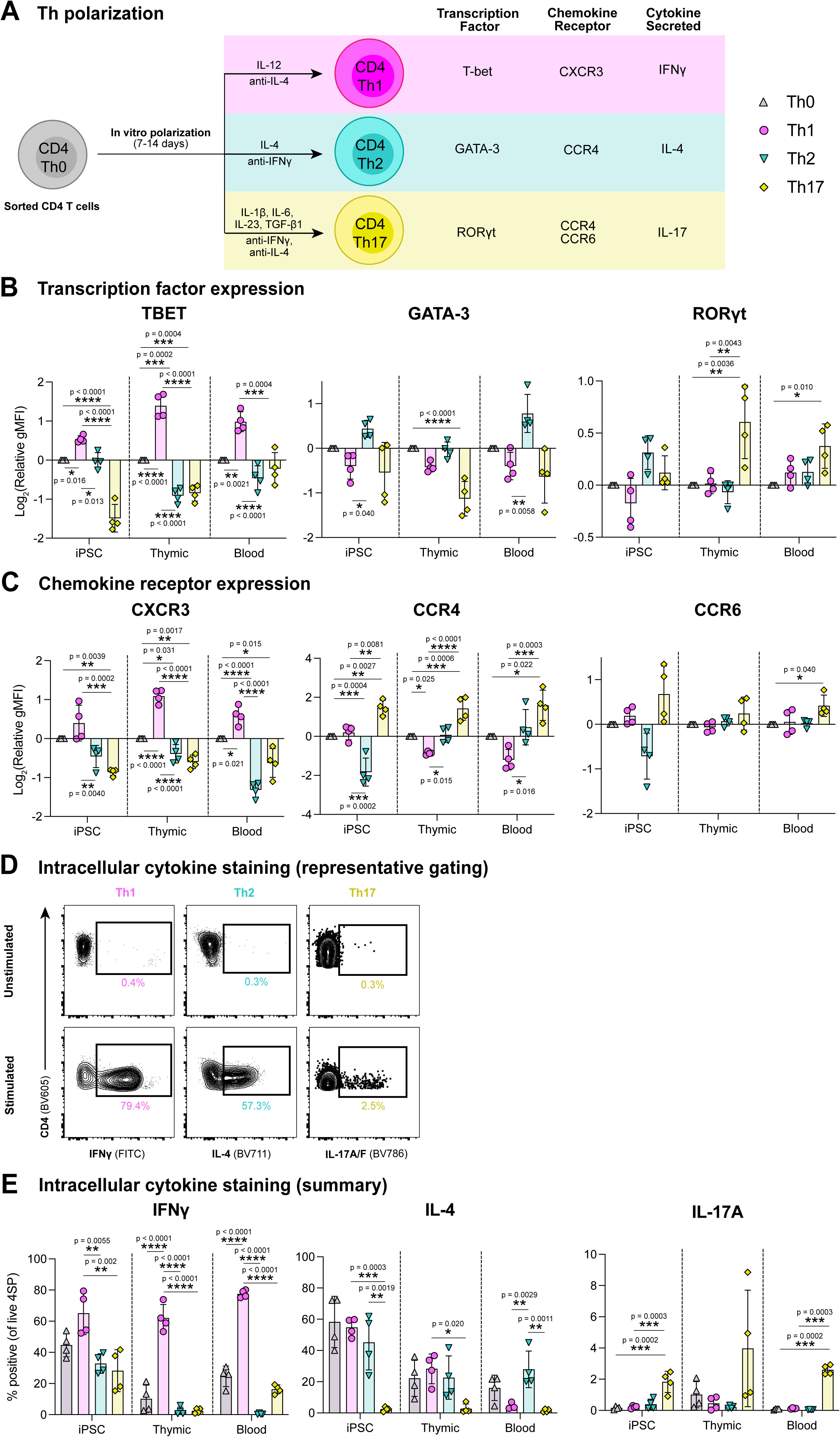
iPSC-CD4^+^ T cells can polarize into functionally different T helper subsets. **(A)** Schematic of polarization and the relevant Th1/2/Th17 transcription factors, chemokine receptors, and cytokines. **(B-C)** iPSC-, thymic-, and blood-CD4^+^ T cells were polarized in Th1/2/17 conditions for 7-14 days, rested overnight in 10 IU/mL IL-2, and then measured for transcription factors (B) and chemokine receptors (C). Data were acquired by flow cytometry and gated on live CD4^+^CD8a^−^. Relative gMFI (geometric mean fluorescent intensity) values normalized to Th0 and then log_2_-transformed. **(D-E)** Rested cells were assessed for intracellular cytokine expression after 4 hours of stimulation with PMA, ionomycin, and Brefeldin A. Data were acquired by flow cytometry and gated on live CD4^+^CD8a^−^. (D) Representative flow plots and (E) quantification of expression of IFNg, IL-4, IL-17A/F in polarized iPSC-CD4^+^ T cells after stimulation with PMA+Iono for 4 hours. Data is shown with mean ± standard deviation; n=4 independent differentiations, or n=4 thymic-or blood-donors collected in 3-4 experiments; statistical significance was determined using ordinary one-way ANOVA with p values shown. See also **Figure S7**.

Polarized cells were phenotyped for Th-related transcription factors and chemokine receptors by flow cytometry and expression levels were normalized to control unpolarized Th0 (**Figures 7B-C** and **S7D-E**). Under Th1 conditions, iPSC-Th1 significantly upregulated TBET expression, and downregulated GATA-3, similar to thymic-and blood-Th1 cells. CXCR3 was upregulated compared to Th2 and Th17 but not Th0 conditions. Under Th2 conditions, iPSC-Th2 cells upregulated GATA-3 similarly to blood-Th2 cells, but downregulated CCR4 unlike thymic-and blood-Th2 cells. Furthermore, while iPSC-Th2s downregulated CXCR3, they did not downregulate TBET. Under Th17 conditions, iPSC-Th17 displayed a trend towards increased RORgt and CCR6 expression that was not statistically significant. However, like thymic-and blood-Th17, iPSC-Th17 cells significantly upregulated CCR4 and downregulated TBET and CXCR3. Overall, these results indicate iPSC-CD4^+^ T cells can be polarized to express Th-specific transcription factors and chemokine receptors.

To determine if polarized iPSC-Th cells can produce hallmark Th cell cytokines, they were stimulated with PMA and ionomycin in the presence of Brefeldin A for 4 hours, then stained for intracellular cytokine expression (**Figures 7D-E**). We found that iPSC-Th1 cells upregulated IFNg expression compared to Th2 and Th17 conditions. Interestingly, neither iPSC-nor thymic-Th1 cells downregulated IL-4 expression, unlike blood-Th1 cells. Under Th2 conditions, IL-4 expression was unchanged for all iPSC-, thymic-, and blood-Th2, but IFNg expression was significantly downregulated relative to their Th1 counterparts. Under Th17 conditions, iPSC-Th17 significantly upregulated IL-17A/F expression relative to Th0, and downregulated IFNg and IL-4 expression relative to their Th1 and Th2 counterparts, with a similar overall pattern of cytokine production compared to thymic-and blood-Th17 cells.

## Discussion

Here we showed that optimization of TCR and Notch signaling enables tunable production of functional CD4^+^ T cells from PSCs. These cells express key markers of naïve T cells, express a diverse TCR repertoire, and can be polarized to Th1-, Th2-, and Th17-like states, producing divergent cytokines. The PSC-CD4^+^ T cell induction protocol works with different fully-defined T cell maturation media formulations and with multiple PSC lines. Collectively, our results have significant potential for advancing allogeneic T cell-based therapies.

It is well known that *in vivo,* longer/stronger TCR stimulation generates CD4^+^ T cells while shorter/weaker stimulation generates CD8^+^ T cells^54^. However, it was unclear how artificial TCR stimulation modalities used for *in vitro* T cell maturation could recapitulate this behavior. Unexpectedly, we found that high levels of some TCR pathway stimulating reagents (anti-CD2/3/28, PHA) bias differentiation towards PSC-CD8^+^ T cells, rather than the expected CD4^+^ T cells. Our data suggest that this is due to a non-monotonic dose-response between the input level of TCR stimulating reagent to the output duration of TCR pathway stimulation: too-low levels lead to insufficient expression of ThPOK and CD8 lineage bias; too-high levels lead to strong TCR downregulation, which may interrupt TCR signaling, similarly to CD8 downregulation following TCR-pMHC Class I interactions^24^. The need to optimize TCR stimulation might explain the lack of PSC-CD4^+^ T cells in many published protocols, even when Notch ligands are not present in the DP-to-SP induction conditions^11,55,56^.

Our study demonstrates that Notch signaling specifically suppresses PSC-CD4^+^ T cell induction. Previous work producing PSC-T cells in artificial thymic organoids (ATOs) found that the use of MS5 feeder cells expressing DLL1 rather than the stronger DLL4 resulted in a higher proportion and yield of mature CD4^+^ T cells, though at the cost of reduced T cell yield overall^30^. These results suggested that weaker Notch ligands may be more permissive to *in vitro* PSC-CD4^+^ T cell induction, though the effect could have been caused by an upstream change in cell development by DLL1 vs DLL4. While still possible, our results show a definitive effect of Notch signaling during *in vitro* positive selection, whereby Notch ligands with increased strength (DLL4>DLL1>JAG2≈JAG1>no ligand) confer greater CD8 lineage bias. Since use of JAG1/2 in ATOs may be too weak to drive PSC-T cell production^37^, appropriately timed DAPT administration during ATO culture could unlock greater PSC-CD4^+^ T cell generation in such systems.

Our data suggest that Notch suppresses the PSC-CD4^+^ T cell development by suppressing TCR signaling, thereby reducing activation of the CD4 lineage-defining transcription factor ThPOK [*ZBTB7B*] and enabling the CD8 lineage-defining transcription factor RUNX3 to dominate and bias differentiation towards CD8^+^ T cells. Interestingly, the role of Notch signaling in *in vivo* DP-to-SP transition is controversial, with studies showing CD4-, CD8-, or no bias following manipulation of Notch signaling^31,32^. Our observation of Notch suppressing TCR stimulation in PSC-DP cells is consistent with one report in NICD-overexpressing mice^57^, but contrasts with most literature showing Notch enhances TCR stimulation^32,58,59^. Thus, the effect of Notch on TCR stimulation is likely context-and dose-dependent^32,57^.

Through comparisons of TCR stimulating reagents, we saw striking differences in the effect of TCR-targeting reagents (anti-CD2/3/28 and PHA) compared to PMA+Iono, which bypasses the TCR to directly activate downstream pathways. Our observation of TCR downregulation immediately after stimulation by anti-CD2/3/28 coincides with data showing anti-CD3 F(ab_2_) fragments cause TCR downregulation and CD8 lineage bias from mouse DP thymocytes^60^. It is thus necessary to fine-tune the dose of these reagents to optimize the duration and intensity of signal. The use of PMA to induce CD4^+^ T cell development traces back to the first studies connecting TCR signaling strength to CD4 lineage commitment in *ex vivo* thymocytes^25,61^. The stronger CD4 bias of PMA+Iono likely stems from avoiding signal interruption by TCR downregulation, though could also result from differential activation of downstream pathways in a manner that favors ThPOK activation^62^. In addition, while PMA gave the highest percent of CD4^+^ T cells, anti-CD2/3/28 and PHA both gave higher yields and expression of maturation markers (e.g. CD27, CD45RA, and CD62L), possibly due to a combination of toxicity by PMA+Iono and positive effects of direct TCR targeting or co-stimulation by the latter reagents. A recent preprint reported PSC-CD4^+^ T cell induction using PMA+Iono at a 10-fold higher PMA dose than used here, applied for 24 hours rather than 1-2 weeks^63^. Consistent with our PMA+Iono results, their PSC-CD4^+^ T cells were predominantly CD45RO^+^CD62L^−^, indicating incomplete maturation.

Through a combination of flow cytometry and scRNAseq, we demonstrated that PSC-CD4^+^ T cells efficiently upregulate naïve T cell markers (such as CD27, CCR7, CD45RA, CD40L, and IL7RA) and CD4 lineage-inducing/specific genes including ThPOK, LEF1, and TCF-1 [*TCF7*]. Importantly, they expressed low levels of cytotoxic genes such as granzyme and perforin. Surface maturation markers Purified PSC-CD4^+^ T cells expanded and upregulated T cell-relevant activation markers in response to stimulation by anti-CD2/3/28. Through *in vitro* Th polarization experiments, PSC-CD4^+^ T cells responded to Th-related cytokine cocktails, expressed relevant transcription factors and chemokine receptors, and secreted relevant cytokines comparably to *ex vivo* thymic-CD4^+^ T cells.

Both PSC-CD4^+^ and CD8^+^ T cells generated diverse TCR repertoires with notable differences in TCR chain usage frequencies compared to thymic-and PBMC T cells. The most striking difference in PSC-T cells was a reduced frequency of Vα and Jα segments at the 5’ end of the V array and 3’ end of the J array, respectively. As these further-apart segments are known to recombine later in SP maturation^50^, PSC-T cells may have an abridged recombination process. Artificial TCR stimulation may trigger positive selection in cells that spent less time recombining the TCRα chain than a typical thymocyte.

Overall, our results are a significant step forward in understanding and controlling PSC-T cell development. The ability to manufacture diverse types of helper T cells opens the door to manufacture therapeutic PSC-derived T cell products for a variety of diseases including cancer, autoimmunity, chronic inflammation, and transplant^1–3^. In particular, our work will enable production of PSC-CAR-T products with defined ratios of CD4^+^ and CD8^+^ T cells, which is expected to improve clinical responses and patient outcomes in the treatment of cancer^16,39^. More broadly, the growing capability to reproduce the blood system from PSCs will enable modeling and exploration of human immune development *in vitro*^64^, production of isogenic cells for immune-competent tissue modeling^65^ and production of cell therapies comprising multiple cooperative immune cell types^66^.

### Limitations of Study

To support robust PSC-T cell production across different experimental runs and from diverse iPSC lines, achieving appropriate purity and developmental stages within CD34^+^ cell populations is critical. This requires carefully optimized media components added at optimal concentrations and times. Additionally, the transition to PSC-CD4^+^ T cell induction relies on precisely timed modulation of Notch and TCR signaling. Since Notch is vital for early T cell development, it must remain active until the population attains a high frequency of DP cells, for which Notch signaling is less essential^67^. This precise timing supports cell population integrity during PSC-CD4^+^ T cell induction. Ongoing improvements in culture conditions, the development of in-process markers for intermediate stages, and the automation of key media perturbations will collectively enhance the robustness and yield of both PSC-CD4^+^ and CD8^+^ T cells. Improvements to CD4^+^ T cell maturation, expansion, and polarization conditions will likewise contribute to improved manufacturing and phenotype stability of diverse helper T cell subtypes for studies testing control of other immune cells and disease states *in vitro* and *in vivo*.

## Resource availability

### Lead contact

Further information and requests for reagents and resources should be directed to and will be fulfilled by the lead contact, Peter W. Zandstra.

### Materials availability

Custom CODEX antibodies are available from the lead contact upon request.

### Data and code availability

- scRNA-seq, scTCR-seq, and CITE-seq data have been deposited at Mendeley Data and will be publicly available as of the date of publication. Accession numbers are listed in the key resources table. Code for data analysis will be available on GitLab as of the date of publication (https://gitlab. com/stemcellbioengineering).
- Any additional information required to reanalyze the data reported in this paper is available from the lead contact upon request.

## Acknowledgements

We thank the thymus donors and their parents for their essential contribution to this research, as well as the surgical and cardiac clinic staff at the British Columbia Children’s Hospital; special thanks to Drs. Sanjiv Gandhi, Andrew Campbell and Al Aklabi, as well as Erin Bleker, Allison Jamieson, Melanie Ganshorn, Aliyah Hassan, Lyn Nguyen and Colleen Ring. We also thank Dr. Sabine Ivison for logistical support in tissue collection. We would like to acknowledge Tara Stach, Yvonne Chung, Stephen Yu, and Bernie Zhao from the BRC-seq core for sequencing support; Michael Williams from the BRC Antibody Lab for technical support and custom reagents; and Andy Johnson and Justin Wong (UBC FACS) and Dr. Lisa Xu and Jessica Huang (BCCHR FACS) for flow cytometry and cell sorting support. We would also like to thank: Dr. Andreas Tiffeau-Meyer for helpful discussion of TCR sequencing analysis; Grace Kuo and Dr. Kelly McNagny for help with annotating innate cell types in scRNAseq data; and Dr. Dominic Boardman and Rosa Garcia for advice on Th polarization experiments. Certain figure cartoons were made with BioRender.

Funding for this work was provided by Genome BC (SIP031 to P.W.Z. and F.M.V.R.) and Wellcome Leap Human Organs, Physiology, and Engineering (HOPE) program (AWD-017859 to P.W.Z., M.K.L., and F.M.V.R.), the Canadian Foundation for Innovation/B.C. Knowledge Development Fund (John Evans Leader’s Fund 38395 to F.M.V.R), and the Canadian Institutes of Health Research (CIHR; ICC-17644 to M.K.L). R.D.J. is supported by a Michael Smith Health Research BC trainee award (RT-2021-1946) and an MSL Pathway to Independence Award. K.S. is supported by CIHR Canadian Graduate Scholarship – Master’s (CGS-M) and Doctoral (CGS-D) award and UBC Faculty of Medicine Graduate Student award. L.N.S is supported by a National Sciences and Engineering Research Council of Canada (NSERC) CGS-D award. F.M.V.R. is supported by CIHR (FDN-159908). M.K.L. is a Canada Research Chair in Engineered Immune Tolerance and receives a Scientist Salary Award from the BC Children’s Hospital Research Institute. P.W.Z. is a Canada Research Chair in Stem Cell Bioengineering.

## Author Contributions

**Conceptualization & Methodology:** R.D.J., K.S., J.M.E., M.K.L., P.W.Z.; **Investigation:** R.D.J., K.S., J.M.E., L.L., J.K.G., A.M., L.J.D., T.P.T., H.H.H., Y.S.M.; **Resources:** T.P.T., H.H.H., C.Z.; **Data curation, Formal Analysis, & Visualization:** R.D.J., K.S., L.N.S., D.R.; **Writing – Original Draft:** R.D.J., K.S., L.N.S.; **Writing – Reviewing & Editing:** R.D.J., K.S., L.N.S., M.K.L., P.W.Z.; **Supervision & Project Administration:** M.K.L., P.W.Z.; **Funding Acquisition:** F.M.V.R., M.K.L., P.W.Z.

## Declaration of Interests

P.W.Z. is a cofounder of Notch Therapeutics. The University of British Columbia has filed a disclosure related to this work on behalf of the authors. The authors declare no other competing interests.

## Supplemental Information

Document S1. Figures S1–S7

Table S1. Intracellular timecourse 3-way ANOVA detailed statistics; related to **Figure 4**.

Table S2. List of genes used in final CD4 and CD8 gene signatures; related to **Figure 5**.

Table S3. Donor information, related to **STAR Methods**.

## STAR METHODS

### Key Resources Table

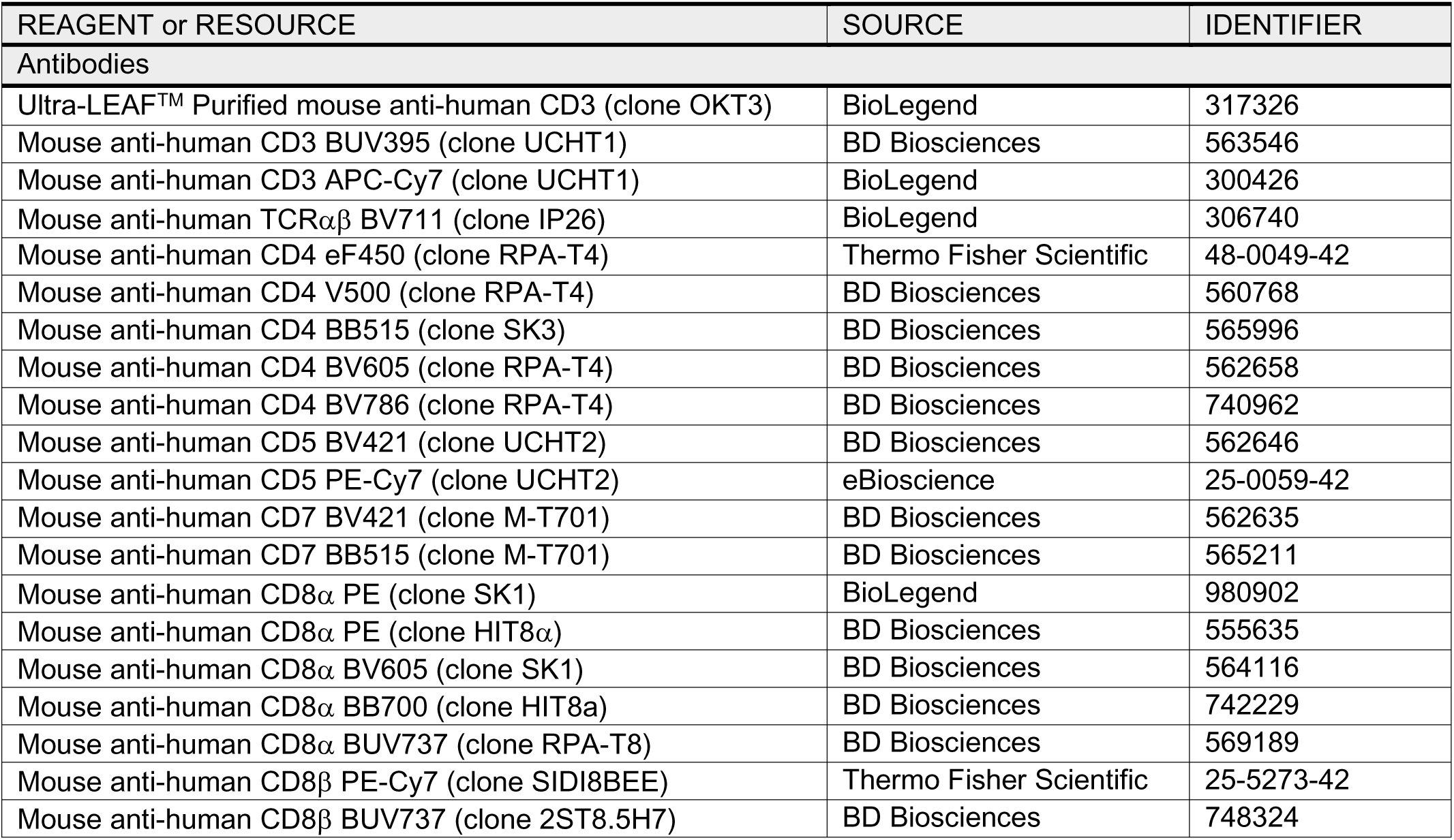
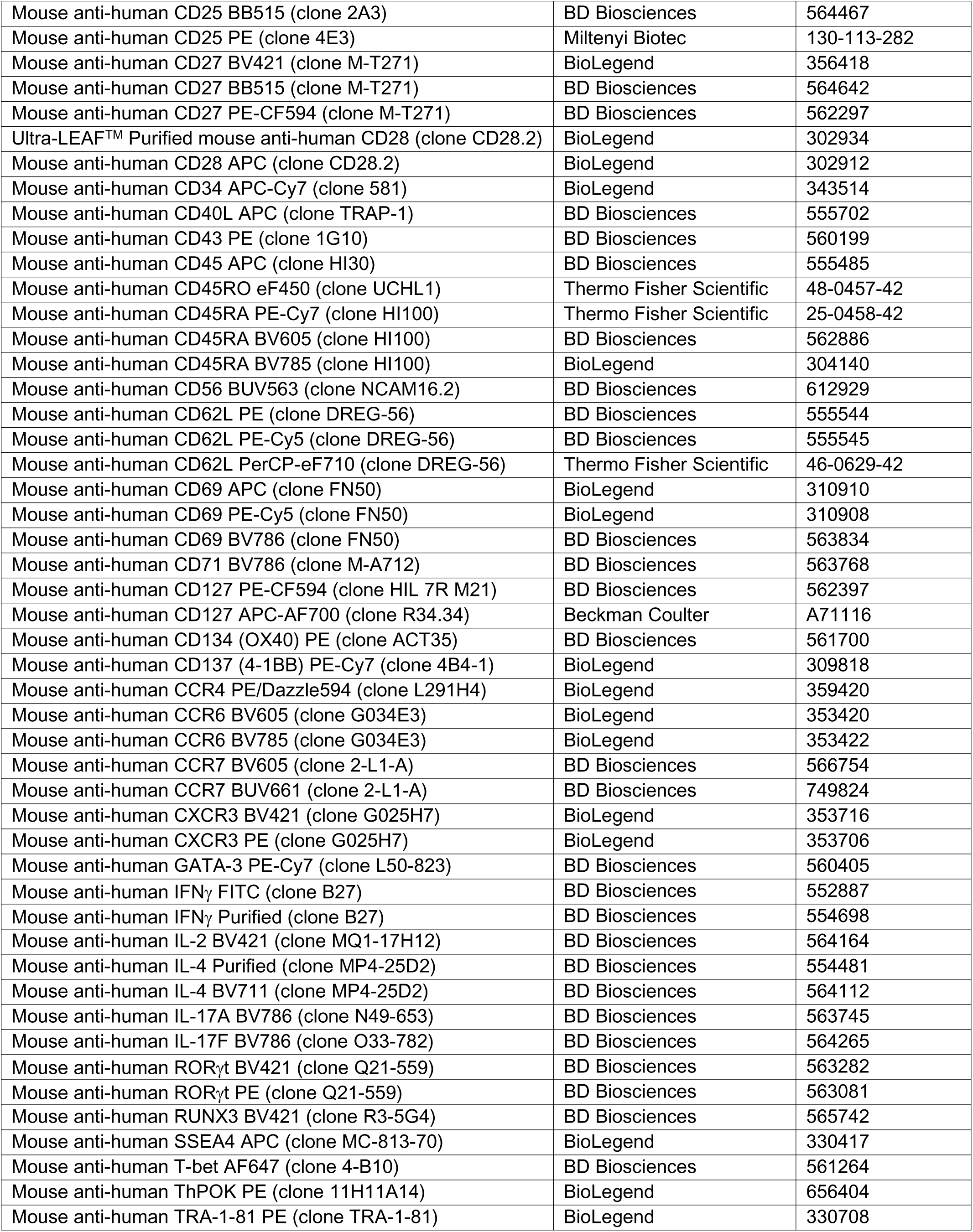
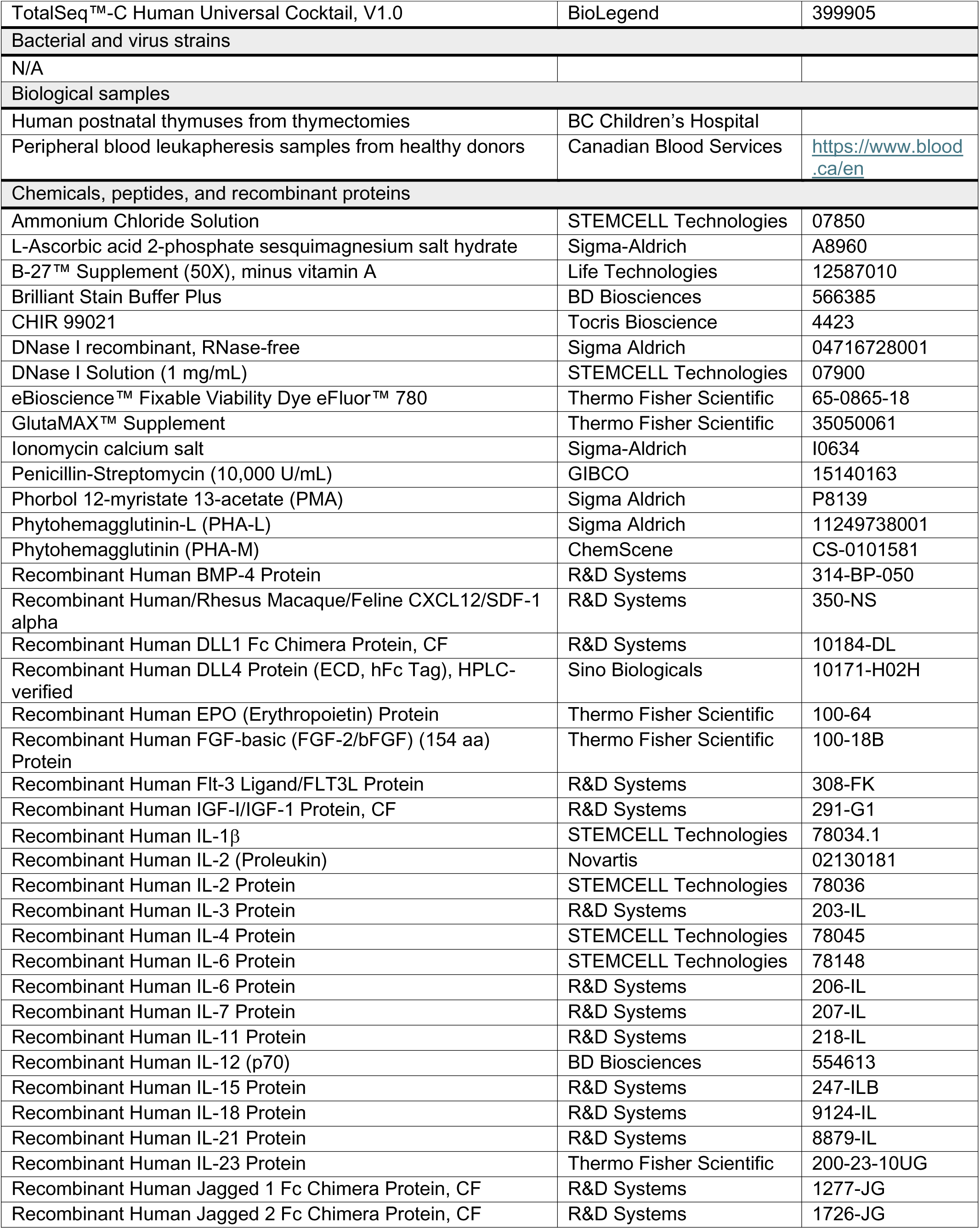
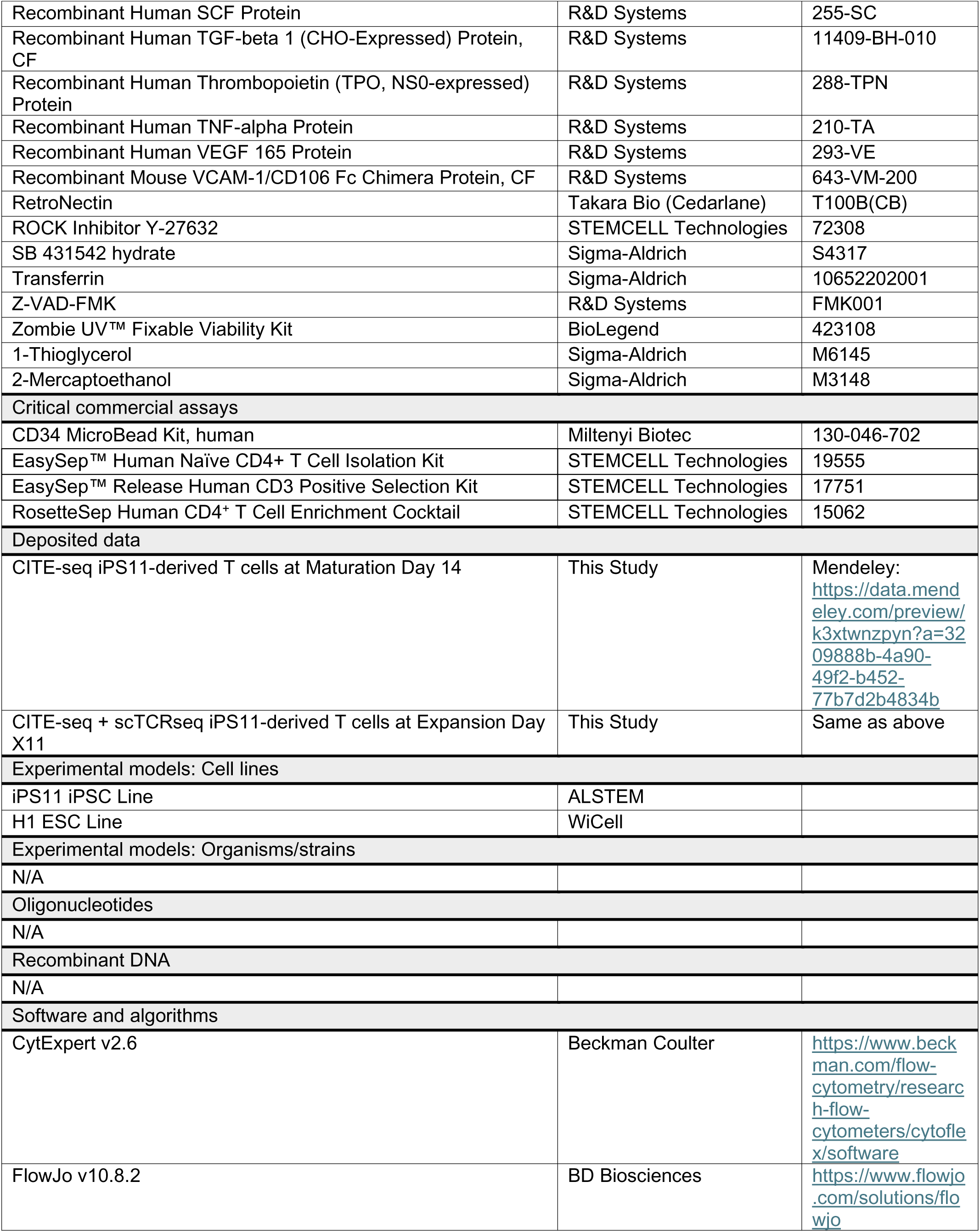
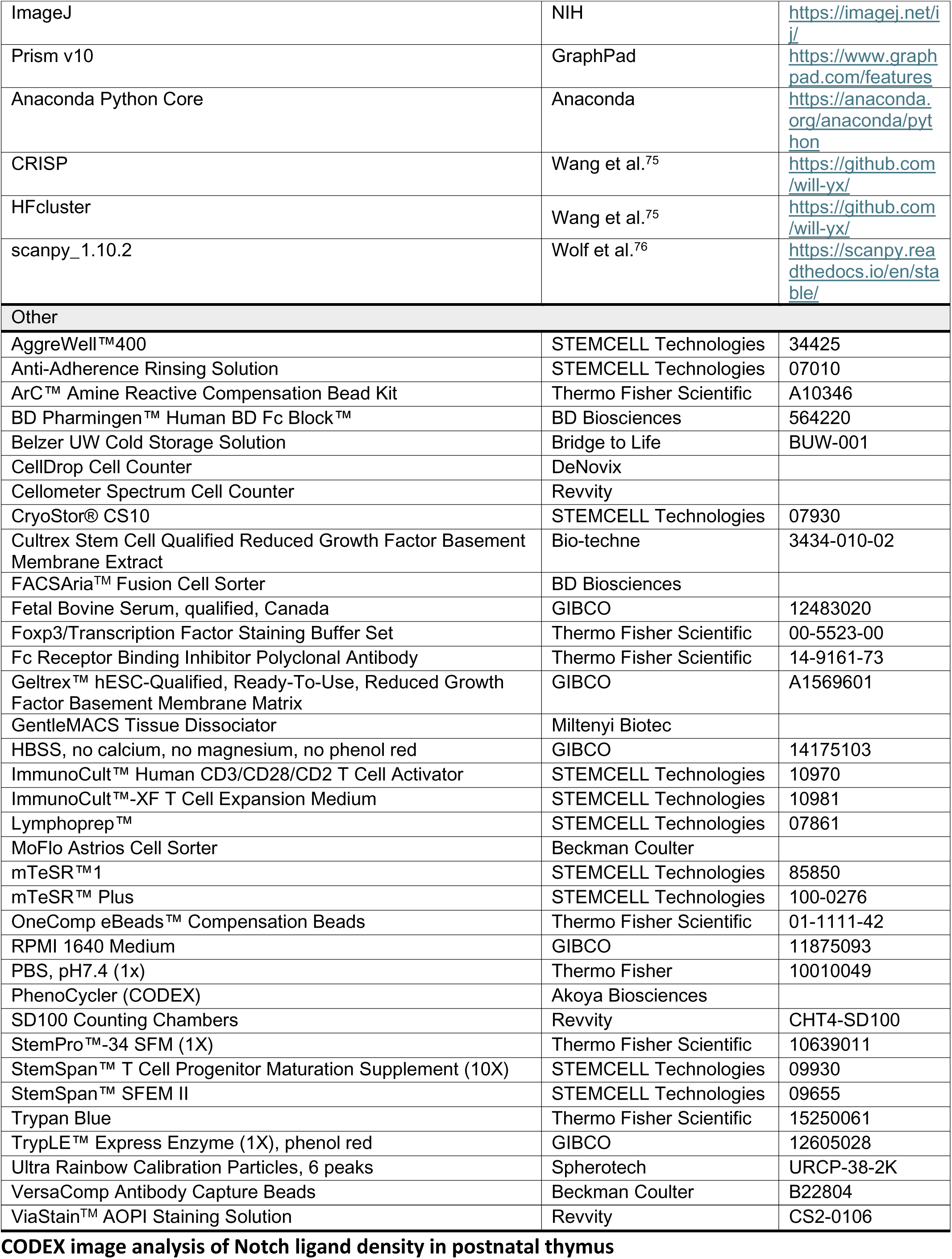

### CODEX image analysis of Notch ligand density in postnatal thymus

Postnatal thymus images were previously acquired via Co-detection by indexing (CODEX) imaging^36^. Three representative images which captured a full thymus lobule (i.e. Capsule, cortex, and full medulla) from donor T085 (M, 8 mo) were chosen for analysis of Notch ligands. For each image, ImageJ was used to draw three lines of thickness 10 from the center of the medulla to the outer edge of the cortex, with the center of the line overlaying the corticomedullary junction. Signal intensity was recorded along the length of the line and binned into 20 segments. The average signal intensity was calculated for each bin. Signal intensities from each line were averaged for each ligand and plotted with GraphPad Prism (v10, GraphPad Software).

### PSC culture

iPS11 cells (ALSTEM, episomal human foreskin fibroblast-derived) and H1 cells^77^ (WiCell), were maintained in mTeSR^TM^1 or mTeSR^TM^ Plus media (STEMCELL Technologies) supplemented with 0.5% penicillin/streptomycin (GIBCO) and in normoxic conditions (20% O_2_, 37°C, 5% CO_2_). Cells were grown on tissue culture-treated plastic plates coated for 1 hour at 37°C with Geltrex (GIBCO) or Cultrex (Bio-Techne). For passaging and seeding HE differentiations, PSCs were dissociated to small clusters by incubating in TrypLE Express (GIBCO) for 2-4 minutes at 37°C, followed by quenching with mTeSR^TM^1 or mTeSR^TM^ Plus and light pipetting. On thaw or passage, cells were treated with 5 μM ROCK inhibitor (ROCKi) Y-27632 (STEMCELL Technologies).

### PSC to HE differentiation

Differentiations of PSCs to CD34^+^ HE cells followed our published AggreWell-based protocol^13^ with minor modifications. AggreWell 400 6-well plates (STEMCELL Technologies) were prepared by coating with 2 mL Anti-Adherence Rinsing Solution (STEMCELL Technologies), spinning down for 5-10 min at 1300 g (to remove bubbles), incubating for 2 hours at room temperature, then washing with 2 mL PBS. PSCs were grown to 50-75% confluency, then dissociated to single cells by incubating in TrypLE Express for 3-5 minutes at 37°C, followed by quenching with mTeSR1 or mTeSR Plus and moderate pipetting. Collected cells were spun down at 200 g for 5 min, then resuspended to 2.21 million cells/mL in “T0” media. T0 media comprises 0.0039% 1-thioglycerol (MTG) (Sigma-Aldrich), 50 μg/mL ascorbic acid (AA) (Sigma-Aldrich), 150 μg/mL Transferrin (Sigma-Aldrich), 10 ng/mL BMP4 (R&D Systems), and 5 μM ROCKi Y-27632 supplemented into StemPro “Complete” base media. StemPro Complete comprises StemPro 34 (Thermo Fisher Scientific) supplemented with 1% Glutamax (Thermo Fisher) and 0.5% Pen/Strep. Aiming for 75 cells/microwell, we added 2 mL T0-cell suspension per 6-well of the AggreWell plate. To achieve uniform aggregation, we allowed cells to evenly settle at RT for 5 minutes, then spun down for 5 minutes at 200 g. Throughout the HE induction, cells were grown in hypoxic conditions (5% O_2_, 37°C, 5% CO_2_).

Differentiating cells were subsequently fed at 24, 42, 72, 96, and 144 hours with “T1”, “T1.75”, “T3”, “T4”, and “T6” media, respectively. The T1, T3, and T6 feeds were each 2 mL media top-ups, while T1.75 and T4 feeds involved carefully aspirating the existing media and adding 2 mL fresh media. Media aspirations and additions were done slowly and at the same edge of each well to minimize disturbances to the aggregates. Each media formulation uses StemPro compete as base and contains the same concentrations of MTG, AA, and Transferrin as in T0 media. T1 media further contains 10 ng/mL BMP4 and 10* ng/mL bFGF (Thermo Fisher) [*the latter achieving a final concentration of 5 ng/mL after topping-up the existing T0 media in each well]. T1.75 and T3 media (equivalent) further contain 10 ng/mL BMP4, 5 ng/mL bFGF, 6 μM SB-431542 (Sigma-Aldrich), and 4 μM CHIR-99021 (Tocris Bioscience). T4 media further contains 5 ng/mL bFGF, 15 ng/mL VEGF (R&D Systems), 10 ng/mL IL6 (R&D Systems), and 5 ng/mL IL11 (R&D Systems). T6 media contains the same factors as T4 media along with 50* ng/mL SCF (R&D Systems), 4* U/mL EPO (Thermo Fisher), and 50* ng/mL IGF-1 (R&D Systems) [*each achieving half the stated concentrations after topping-up the existing T4 media in each well].

CD34^+^ HE cells were collected 192 hours (day 8, “T8”) after initiation of differentiation. Aggregates were collected, spun down at 200 g for 5 min, then dissociated to single cells by incubating in 0.5 mL (per pooled 6-well) TrypLE supplemented with 100 U/mL DNase I (Sigma-Aldrich) for 15 minutes at 37°C, vigorously pipetting every 5 minutes. The TrypLE was quenched with a 50:50 mix of HBSS (GIBCO) and FBS (GIBCO). Cells were spun down for 5 min at 200 g, washed in 3 mL HBSS +2% FBS, then resuspended in 1 mL HBSS +2% FBS for pre-enrichment counting; a subset of cells was set aside for flow cytometry. CD34^+^ cells were isolated using the human CD34-MicroBead kit (Miltenyi Biotec), as per the manufacturer’s instructions. Post-enrichment cells were counted and frozen down in CryoStor CS10 (STEMCELL Technologies) at 1 million cells/mL; a subset of cells was set aside for flow cytometry. The set-aside pre-and post-enrichment cells were stained with antibodies against key markers: CD34, CD43, CD73, and CD184 to validate HE induction^78^.

### PSC-HE to T cell differentiation

PSC-T cell differentiations generally followed our published protocol^13^, with minor modifications. In all cases, cells were incubated in normoxic conditions (20% O_2_, 37°C, 5% CO_2_). For PSC-HE to HSPC induction, wells were prepared by coating TC-treated plates with 10-15 (typically 15) μg/mL hDLL4-Fc (Sino Biological) + 2.5 μg/mL mVCAM1 (R&D Systems) for 2 hours at RT or 37°C, or overnight at 4°C. PSC-HE cells were thawed and seeded at 30-100 (typically 100) x 10^3^ cells/mL (3-10 x 10^3^ cells/100 μL per 96 well) in EHT media. EHT media contains the same factors as T4 media (see above) with VEGF reduced to 5 ng/mL and additionally supplemented with 10 ng/mL BMP4, 50 ng/mL SCF, 30 ng/mL TPO (R&D Systems), 25 ng/mL IGF-1, 10 ng/mL IL3 (R&D Systems), 10 ng/mL Flt3L (R&D Systems), and 10 μM ROCKi Y-27632. Cells were passaged to Pro-T induction conditions at EHT day 4-7 (annotated E4-7; typically E5).

For PSC-HSPC to Pro-T induction, wells were prepared by coating with 10-15 (typically 15) μg/mL hDLL4-Fc + 2.5 μg/mL mVCAM1 for 2 hours at RT or 37°C, or overnight at 4°C. Cells output from the HSPC induction stage were seeded at 30-100 (typically 100) x 10^3^ cells/mL (3-10 x 10^3^ cells/100 μL per 96 well) in “PSC1” media. PSC1 comprises 12.37 ng/mL SCF, 8.61 ng/mL Flt3L, 97.4 ng/mL CXCL12 (R&D Systems), 0.07 ng/mL TNFα (R&D Systems), 0.97 ng/mL IL3, and 65.25 ng/mL IL7 (R&D Systems) supplemented into “JAC Ultra” base media. JAC Ultra comprises IMDM with GlutaMAX (GIBCO) supplemented with 4% B27 without Vitamin A (Thermo Fisher), 0.5% Pen/Strep, 24 μM BME (Sigma-Aldrich) and 60 μM AA. Cells were fed by top-up with an equal volume of fresh media at Pro-T induction day 3 or 4 (annotated P3 or P4). Cells were sampled for flow cytometry and passaged to DP induction conditions between day P7-9 (typically P7).

For iPSC-Pro-T to DP maturation, wells were prepared by coating with 10-15 (typically 10) μg/mL hDLL4-Fc + 2.5 μg/mL mVCAM1 for 2 hours at RT or 37°C, or overnight at 4°C. Cells output from the Pro-T induction stage were seeded at 0.5-4 (typically 2) x 10^6^ cells/mL (50-400 x 10^3^ cells/100 μL per 96 well) in “PSC2” media. PSC2 comprises 9.76 ng/mL SCF, 4.96 ng/mL Flt3L, 15.22 ng/mL CXCL12/SDF-1, 0.04 ng/mL TNFα, 2.55 ng/mL IL3, and 71.93 ng/mL IL7 supplemented into “JAC Ultra” base media. Where indicated, cells were instead seeded into media comprising StemSpan T cell Progenitor Maturation Supplement (STEMCELL Technologies) diluted 1:10 into either JAC Ultra or SFEM II (STEMCELL Technologies). Cells were fed by top-up with an equal volume of fresh PSC2 media at DP maturation day 3 or 4 (annotated M3 or M4). Thereafter, cells were fed by 50% media exchange with an equal volume of fresh media every 3-4 days. Cells were typically sampled for flow cytometry every ∼7 days (e.g. days M7, M14, and M21). Cells were induced to SP cells starting between day M14-28 (typically M14 or M21), aiming for the population of CD3^+^TCRαβ^+^ cells to be >10% among live cells.

For iPSC-DP to SP induction, wells were prepared by coating with 0-10 μg/mL hDLL4-Fc, hDLL1-Fc (R&D Systems), hJAG1-Fc (R&D Systems), or hJAG2-Fc (R&D Systems) + 2.5 ug/mL mVCAM-1 for 2 hours at RT or 37°C, or overnight at 4°C. Where indicated, the wells were left uncoated. Cells output from the DP induction stage were seeded 1:1 in PSC2 supplemented with 0-2.5% Immunocult anti-CD2/3/28 complexes (STEMCELL Technologies), 0-2.5 μg/mL PHA-M (**Figure 2**, **Figure S3**, ChemScene) or PMA-L (**Figure 3**, **Figure S4**), or 0-0.25 ng/mL PMA (Sigma-Aldrich) + 100 ng/mL Ionomycin (Sigma-Aldrich). Where indicated, cells were instead seeded into StemSpan maturation supplement + JAC Ultra or SFEM II. Where indicated, the cells were stimulated by 50-90% media exchange with fresh media containing stimulating reagents. Cells were fed by top-up with an equal volume of fresh media (without stimulating reagents) at SP induction day 3 or 4 (annotated S3 or S4). Cells were transferred to expansion conditions after 7-14 days. For 14 day stimulations, cells were either passaged 1:1 onto wells freshly coated with 2.5 μg/mL mVCAM1 only, or fed by 50% media exchange, both with fresh media without stimulation, as indicated in figure legends. In both cases, cells were fed again with fresh media (without stimulation) at day S10 or S11. Cells were typically sampled for flow cytometry on days S7 and S14. In some cases, cells were sampled for flow cytometry at earlier timepoints (e.g. 24-72 hours after stimulation).

### PSC-CD4^+^ T cell generation and isolation for expansion and polarization tests

iPSC-HE to HSPC induction and iPSC-HSPC to Pro-T induction were carried out as above. For iPSC-Pro-T to DP maturation, wells were prepared by coating with 15 μg/mL hDLL4 + 2.5 μg/mL mVCAM1. iPSC-Pro-T cells were seeded at 3 x 10^6^ cells/mL (300 x 10^3^ cells/100 μL per 96 well) in StemSpan media. (StemSpan T cell Progenitor Maturation Supplement diluted 1:10 into SFEM II; STEMCELL Technologies). Cells were fed by top-up with an equal volume of fresh StemSpan media at DP maturation day M3 or M4, then fed by 50% media exchanges every 3-4 days until day M14. For iPSC-DP to 4SP induction, day M14 cells were collected, centrifuged, resuspended in StemSpan media supplemented with 0.5 μg/mL PHA-L for stimulation, and plated into uncoated 6 well plates (9 x 10^3^ cells/5mL per 6 well). Cells were fed by 50% media exchange with an equal volume of fresh StemSpan media (without stimulation) 3-4 days post-stimulation (S3-4). On days S7 and S10, cells were fed by 50% media exchange with Immunocult-XF T Cell Expansion Media (STEMCELL Technologies) supplemented with 200 IU/mL IL-2 (Proleukin). On day S14, cells were collected, stained for antibodies for 15 min at room temperature, and sorted for iPSC-CD4^+^ T cells (TCRab^+^CD4^+^CD8a^-^) using MoFlo Astrios (Beckman Coulter) or FACSAria Fusion (BD Biosciences). Flow-sorted cells were subsequently used for experiments.

### CD4^+^ T cell isolation from human postnatal thymus

Human postnatal thymus tissue was collected in UW Solution (Bridge to Life) with consent from infants (donor sex and age can be found in **Table S3**) undergoing cardiac surgeries in BC Children’s Hospital. A piece of the thymus tissue was mechanically dissociated in Immunocult-XF T cell Expansion Medium using GentleMACS Dissociator (Miltenyi Biotec). The dissociated bulk thymocytes were filtered and subsequently frozen for further use. On day of experiment, frozen bulk thymocytes were thawed, treated with DNAseI (1 mg/mL final concentration; STEMCELL Technologies) for 10 min at room temperature, stained with antibodies at 15 min at room temperature, and flow-sorted for thymic-CD4^+^ T cells (TCRab^+^CD25^-^CD4^+^CD8a^-^) using FACSAria Fusion (BD Biosciences). Flow-sorted cells were subsequently used for experiments.

### CD4^+^ T cell isolation from human adult peripheral blood

Leukapheresis products were obtained from consented healthy adult donors and enriched for CD4^+^ cells prior to flow sorting (donor sex and age can be found in **Table S3**). Blood samples were incubated with RosetteSep Human CD4^+^ T Cell Enrichment (STEMCELL Technologies) at room temperature for 20 minutes, diluted in 1:1 ratio with 1X PBS (GIBCO), layered atop Lymphoprep (15 mL/tube, STEMCELL Technologies), and fractionated by centrifugation (582 x g, 25 min, no brake, room temperature). The buffy coat layer was collected using transfer pipettes, and red blood cells within the layer were lysed using Ammonium Chloride Solution (5 mL/donor, STEMCELL Technologies) for 5 min at room temperature. Platelets were then removed by centrifugation (129 x g, 10 min, room temperature). The purified CD4^+^ cells were minimally depleted of CD45RO^+^ cells using half of the recommended concentration of EasySep Human Naïve CD4^+^ T Cell Isolation Kit and Magnet (STEMCELL Technologies). Negative fraction from the magnetic isolation were stained with antibodies for 15 min at room temperature, and flow-sorted for naïve blood CD4^+^ T cells (CD4^+^CD25^-^CD127^+^CD45RA^hi^CD45RO^-^CD62L^hi^) using FACSAria Fusion (BD Biosciences). Flow-sorted cells were frozen and thawed on the day of experiment.

### T cell expansion and polarization

For all polarization and expansion experiments, cells were incubated in normoxic conditions (20% O_2_, 37°C, 5% CO_2_), maintained at a concentration of 0.5 x 10^6^ cells/mL and cell density of 0.3 x 10^6^ cells/m^2^, and expanded in Immunocult-XF T Cell Expansion Media (STEMCELL Technologies) supplemented with 1X Pen/Strep (GIBCO), 100 IU/mL IL-2 (Proleukin). For polarization experiments, the cells were further supplemented with the following Th1/Th2/Th17 polarization cocktails for the entirety of culture period. Th1 cytokine cocktail: 10 ng/mL IL-12, 1 μg/mL anti-IL-4; Th2 cytokine cocktail: 10 ng/mL IL-4, 1 μg/mL anti-IFNg; and Th17 cytokine cocktail: 10 ng/mL IL-1b, 10 ng/mL IL-6, 20 ng/mL IL-23, 10 ng/mL TGF-b1, 5 μg/mL anti-IFNg, and 5 μg/mL anti-IL-4.

On day 0, iPSC-, thymic-, or blood-CD4^+^ T cells were stimulated with 2.5% anti-CD2/3/28 (STEMCELL Technologies). Cells were split every 2-3 days and replated at the above cell density/concentration, with cytokines replenished. iPSC-and thymic-CD4^+^ T cells were restimulated on day 7 with 2.5% anti-CD2/3/28, and undergo an additional 7 days of expansion/polarization. On day 14 (iPSC-and thymic-CD4^+^ T cells) or day 7 (blood-CD4^+^ T cells), cells were collected, washed, and resuspended in fresh Immunocult-XF T Cell Expansion Media supplemented with only 10 IU/mL IL-2 (Proleukin), and rested for 16-24 hours prior to running downstream assays. Post-resting, cells were sampled and phenotyped on FACSymphony A5 (BD Biosciences).

### T cell activation assay and intracellular cytokine staining

For activation assays, post-rested cells were washed twice with PBS, and plated at 100 x 10^3^ cells/200 μL in a 96 well flat-bottom, in Immunocult-XF T Cell Expansion Media supplemented with 10 IU/mL IL-2 and 2.5% anti-CD2/3/28 complexes (STEMCELL Technologies). Cells were incubated in normoxic conditions (5% O_2_, 37°C, 5% CO_2_) for 48 hours. Post-stimulation, cells were harvested, stained, and phenotyped for activation markers on FACSymphony A5 (BD Biosciences).

For intracellular cytokine secretion assays, post-rested cells were washed twice with PBS, and plated at 50 x 10^3^ cells/200 μL in a 96 well round-bottom, in Immunocult-XF T Cell Expansion Media. Unstimulated cells were supplemented with 10 μg/mL Brefeldin A. Stimulated cells were supplemented with 10 μg/mL Brefeldin A, 10 ng/mL PMA, and 500 ng/mL ionomycin. Cells were incubated in normoxic conditions (5% O_2_, 37°C, 5% CO_2_) for 4 hours. Cells were then harvested, stained, and phenotyped on FACSymphony A5 (BD Biosciences).

### Flow cytometry

Antibodies can be found in **Key Resources Table**. HBSS or PBS supplemented with 2% FBS was used as Flow buffer. Staining was done entirely in 96 well V-bottom plates. Cells were collected and washed once with PBS. Cells were stained with Fc-blocking antibodies (BD Biosciences or Thermo Fisher) to reduce non-specific binding, Fixable Viability Dye (BioLegend or Thermo Fisher) to exclude dead cells. Cells were stained for surface proteins in PBS or Flow buffer for 30 min in the dark at 4°C or room temperature. Brilliant Plus Buffer was included in staining mixes when using at least three BV or BUV antibodies. Cells were washed with PBS or Flow buffer, resuspended in Flow buffer, and acquired on FACSymphony A5 (BD Biosciences) or CytoFLEX LX N3-V5-B3-Y5-R3-I0 (Beckman Coulter).

For detection of intracellular proteins, cells were washed with PBS of Flow buffer, fixed and permeabilized with eBioscience Foxp3/Transcription Factor Staining Buffer Set (Thermo Fisher) for 45 min in the dark at room temperature or 4°C overnight, then stained for intracellular proteins for 45 min in the dark at 4°C or room temperature. Brilliant Plus Buffer was included in intracellular staining mixes when using at least three BV or BUV antibodies. Cells were washed and resuspended in Perm Buffer from the eBiosciences kit and acquired on the FACSymphony or CytoFLEX.

Data were analyzed on FlowJo software (v10.X; BD Biosciences) or CytExpert (v2.6; Beckman Coulter), then exported for plotting and statistical analysis with GraphPad Prism (v10, GraphPad Software) or Python (v3.11.4).

### Three-way ANOVA for intracellular staining timecourse

The Python package statsmodels (v0.14.0) was used for three-way ANOVA modeling. Each population percent or log-transformed geomean value was fit with an ordinary least squares model of the form:

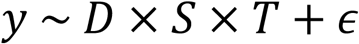

Where *y* is the measured value and is modeled as a full-factorial function of *D*: unique combinations of DLL4 and DAPT inputs (−DLL4−DAPT, +DLL4−DAPT, +DLL4+DAPT), *S*: levels of anti-CD2/3/28 (0.3%, 1.25%), *T*: time, and *ϵ*: the residual error. Hypotheses were tested with Type II sum-of-squares. A threshold of p = 0.01 was considered significant. Detailed statistics are provided in **Table S1**.

### Sample preparation for scRNAseq and CITE-seq

The BioLegend TotalSeq-C CITE-seq antibody preparation and 10X Genomics CITE-seq cell staining protocols were followed. Briefly, one vial of TotalSeq-C Human Universal Antibody Cocktail v1.0 (BioLegend) was equilibrated to room temperature for 5 min and then spun at 10,000g for 30 seconds. The lyophilized panel was resuspended in 27.5uL of HBSS + 4% FBS, vortexed for 10 seconds, and then incubated at room temperature for 5 min. The vial was vortexed again for 10 seconds and then spun at 10,000g for 30 seconds. The entire volume was transferred to a low protein binding PCR tube and then centrifuged at 14,000g for 10min at 4°C.

Prior to CITE-seq antibody staining, 250,000 total cells were partitioned into a 12 x 75mm tube. Cells were spun at 500xg for 5 min at 4°C and resuspended directly in 12.5uL of HBSS + 4% FBS. 12.5uL of the antibody staining cocktail was added and the cells were incubated for 30min at 4°C. Cells were washed three times with 3.5mL HBSS + 4% FBS and resuspended in a final volume of 55uL. Cells were counted, stored on ice, and given immediately to the sequencing facility for 10X Genomics 5’ library preparation and sequencing.

The generation of single cell indexed libraries was performed by the Biomedical Research Center Next Generation Sequencing Core using the 10X Genomics Chromium Controller platform and the Chromium Single Cell 5’ Library and Gel Bead Kit v1.1 and Chromium Single Cell Library and Gel Bead Kit v2 reagents. The sequencing protocol provided by the supplier was followed without modification for CITE-seq and run with the NextSeq 2000. After run completion, the Binary base call (bcl) files were converted to fastq format using the Illumina bcl2fastq2 software, and data were received for further analysis.

### scRNAseq and CITE-seq data analysis

Raw fastq files from all samples were aligned and quantified using CellRanger (CellRanger 6.0.1) and aligned with the human genome reference hg38. Following the suggested pipeline for quality control in Scanpy (v 1.10.2), CITE-seq data were filtered for dead cells, doublets, and red blood cells by excluding cells with greater than 5% mitochondrial genes and less than 1000, but no more than 30000 genes. Genes found in less than 3 cells were removed. Samples underwent normalization, scaling, dimensional reduction, and further downstream analysis using the standard Scanpy workflow. Leiden clustering was performed using 50 principal components and a resolution of 0.5. Cells were clustered and identified based on known marker genes.

### CD4 vs CD8 gene signature analysis

Using two distinct human T cell development single-cell RNA sequencing datasets (Chopp *et al*.^47^ and Park *et al*.^50^), we generated gene signatures of thymic CD4^+^ and CD8^+^ T cells to measure the likeness of iPSC-derived populations. The top 20 marker genes for each relevant CD4^+^ or CD8^+^ T cell cluster were aggregated, then filtered to remove duplicate genes. Chopp *et al*. CD4 clusters: “hs-ImCD4”, “hs-MatCD4”; Park *et al*. CD4 clusters: “CD4+T”, “CD4+Tmem”; Chopp *et al*. CD8 clusters: “hs-ImCD8”, “hs-MatCD8”, “hs-CD8”; Park *et al*. CD8 clusters: “CD8+T”, “CD8+Tmem”. By taking the union of genes across these CD4-or CD8-related clusters, we aimed to ensure our signatures were robust to manual annotation and batch effects. Finally, genes present in both the CD4 and CD8 signatures were excluded. The final CD4 and CD8 signatures contain 49 and 63 genes, respectively (see **Table S2**). Signature scores were generated with Scanpy’s gene set scoring function (scanpy.tl.score_genes), which normalizes the average z-score of signature gene expression against a set of 50 genes in a standard reference set.

### TCR repertoire analysis

Signac (v0.17.0) was used to process scTCR-seq data. iPSC-T cell scTCRseq data were compiled with fetal and postnatal thymocyte data^50^ and PBMC data^51^. As an unbiased estimator of TCR repertoire diversity, we used Simpson’s Diversity (*D*). This was computed on TCR clones (unique productive rearrangements of CDR3, V, and J segments) separately for TCRα and β chains using the formula:

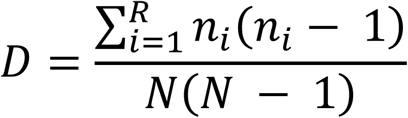

where *R* is the total number of TCR clones, *n*_*i*_ is the number of templates for each clone *i*, and *N* is the total number of TCR templates. The value *D* is interpreted as the probability that two sampled templates are from the same TCR clone^79^. *D* = 0 indicates each sequencing template was sampled from a unique clone, while *D* = 1 indicates all templates were sampled from the same clone. In postnatal thymocyte samples that we evaluated^50^, the number of cells was in the 100’s and no templates came from the same clone.

The variance (*D*_*var*_) was approximated using a higher moment^71^:

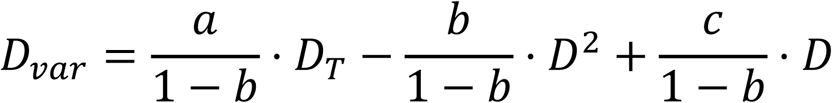

Where

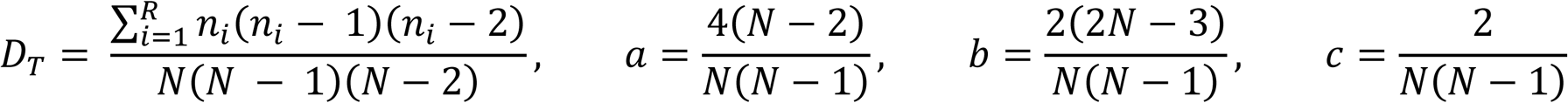

For plotting, we show *D* ± *D*_*σ*_, with the standard deviation estimated as 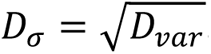.

Segment usage densities were smoothed with a B-spline representation, with a uniform weight of 1/*σ* applied (*σ* = standard deviation of segment usage frequencies), and smoothing factor *s* given by the number of segments per V or J array.

---

**Figure S1.**
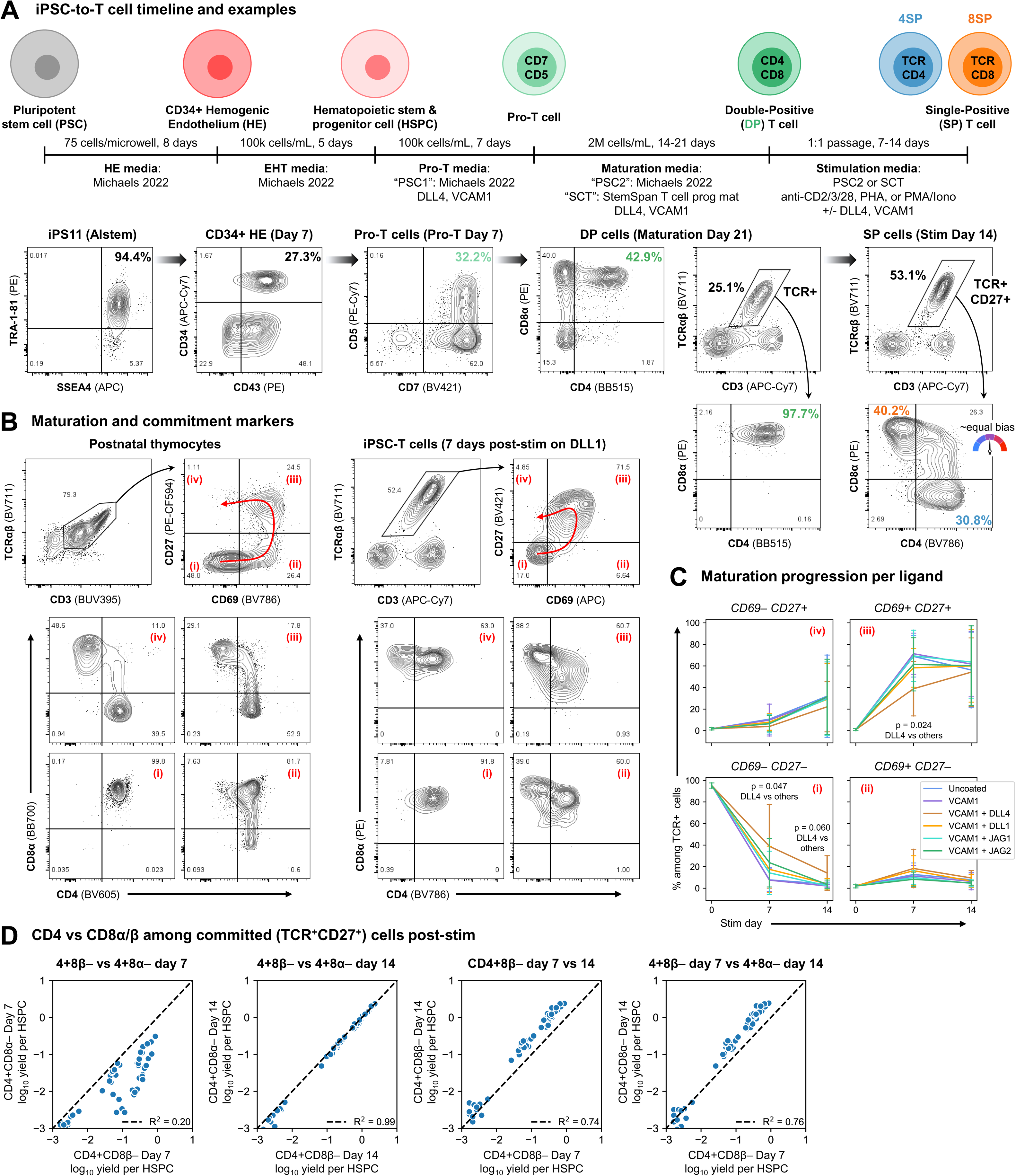
Details of iPSC-to-T cell differentiation, related to Figure 1. **(A)** Media, timelines, and representative flow plots showing key cell types during iPSC-to-T cell development. iPSCs are aggregated, then induced into mesoderm and arterial hemogenic endothelium (HE)^13^. CD34^+^ iPSC-HE are induced to generate budded CD34^+^CD43^+^CD45^+^ iPSC-HSPCs via an endothelial-to-hematopoietic transition (EHT)^13^. iPSC-HSPCs are induced towards CD5^+^CD7^+^ iPSC-Pro-T cells, then iPSC-DP cells in a two-part serum-and feeder-free engineered thymic niche (ETN) comprising immobilized DLL4 and VCAM1 proteins and two staged formulations of soluble cytokines including IL7, Flt3L, CXCL12 (SDF-1), IL3, and SCF, referred to as “PSC1” (phase 1) and “PSC2” (phase 2)^13,72,73^. The ∼equal bias 4SP/8SP sample is two weeks after inducing to SPs with 0.5% anti-CD2/3/28 on wells coated with 2.5 μg/mL VCAM1 only (no DLL4). **(B)** Progression of cells from unsignaled DP cells to mature, committed SP cells. To track the progress of positive selection and subsequent maturation, we use the TCR stimulation marker CD69 and lineage commitment marker CD27^35^ to classify: (i) CD69^−^CD27^−^ immature “unsignaled” cells; (ii) CD69^+^CD27^−^ “signaled” but uncommitted cells; (iii) CD69^+^CD27^+^ recently-committed cells; and (iv) CD69^−^CD27^+^ mature, committed cells. Left: postnatal thymocytes, right: iPSC-derived T cells (DLL1+DMSO condition from Figure 1F). The top plots show the gating of TCR^+^ cells into the CD69/CD27 populations, the bottom plots show the CD4 vs CD8 distribution within each population. **(C)** CD69 vs CD27 expression dynamics on cells stimulated in the presence or absence of different Notch ligands from Figure 1F. n=4 independent differentiations; errorbars: mean ± standard deviation; Statistical significance was determined using unpaired T-tests between DLL4 and the other conditions pooled together at stim day 7 or 14, with p values shown. **(D)** Faster downregulation of CD8β than CD8α in committed CD27^+^ T cells during iPSC-DP-to-SP induction. Correlation between yields per CD34^+^CD43^+^ HSPCs of CD3^+^TCRαβ^+^CD27^+^ 4SPs that are CD4^+^CD8α^−^ or CD4^+^CD8β^−^ after 7 or 14 days of DP-to-SP induction. Collated from samples in Figure 1E-F.

**Figure S2.**
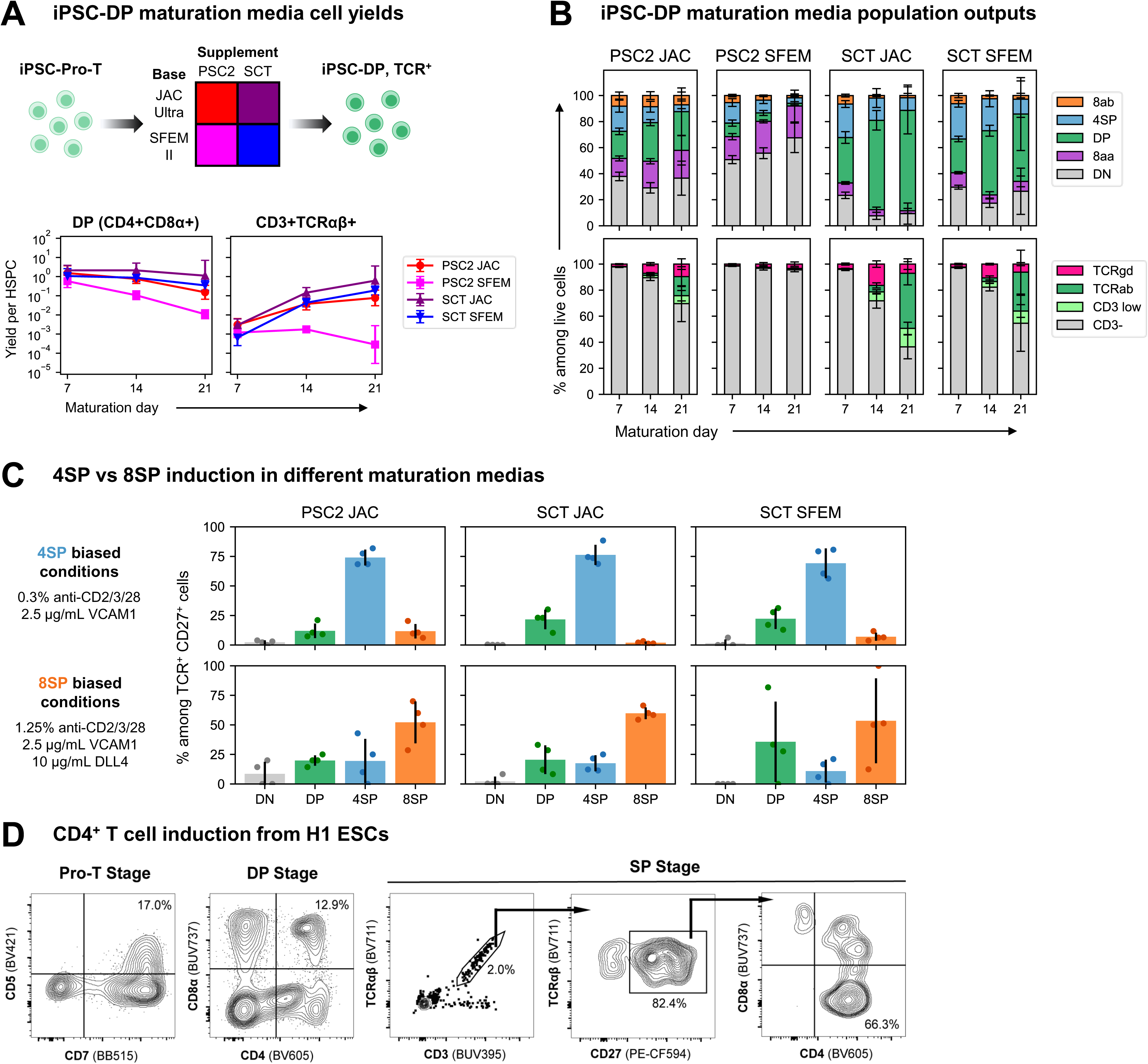
PSC-CD4^+^ T cell induction is robust to choice of maturation media and cell source, related to Figure 1. **(A)** iPSC-Pro-T cells were differentiated to DP and TCR^+^ cells in four maturation media variants. Each maturation media was a combination of a cytokine supplement from either our prior work (PSC2^13^) or StemCell Technologies (SCT, T cell Progenitor Maturation Supplement) added to a base media from either our prior work (JAC Ultra^13^) or StemCell (SFEM II). Yields of DP and TCR^+^ cells per CD34^+^CD43^+^ HSPC input were measured during 3 weeks of maturation in wells coated with 10 μg/mL DLL4 + 2.5 μg/mL VCAM1. n=4 independent differentiations; errorbars: mean ± standard deviation. **(B)** Comparison of population composition during in vitro iPSC-DP/TCR+ cell maturation in different media variants. 4SP cells here are nearly all TCR^−^ CD4 immature single positive cells (4ISPs). **(C)** Comparison of DP vs SP phenotype distributions following DP-to-SP induction in CD4-or CD8-biased conditions. Cells were induced to SPs in the same media used for maturation. 2D cell input; n=4 independent differentiations; errorbars: mean ± standard deviation. **(D)** Confirmation that iPSC-T cell production and iPSC-CD4+ T cell induction conditions work for H1 ESCs (WiCell). H1 cells were differentiated to PSC-Pro-T cells using our published protocol^13^, then transferred to SCT media for PSC-DP cell maturation. Cells were induced to 4SP by stimulating with 0.5 μg/mL phytohemagglutinin (PHA, see Figure 2) in uncoated wells.

**Figure S3.**
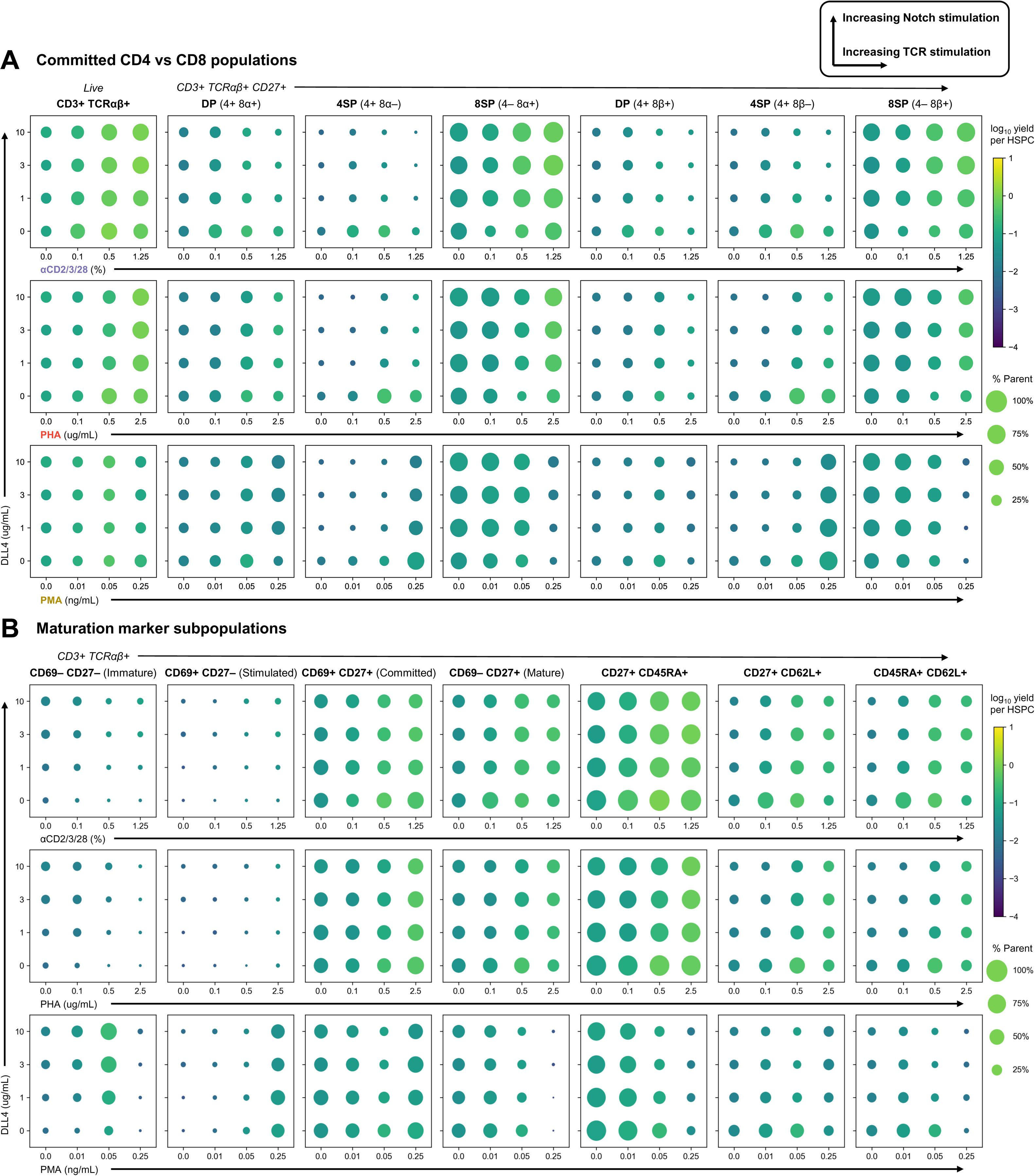
Additional data for TCR stimulation reagents, related to Figure 2. **(A, B)** Populations of interest generated in each combination of Notch and TCR stimulating reagent level. The size and color of each dot represents the percent of each population and the yield per CD34^+^CD43^+^ HSPC input, respectively. Values are mean of n=4 independent differentiations.

**Figure S4.**
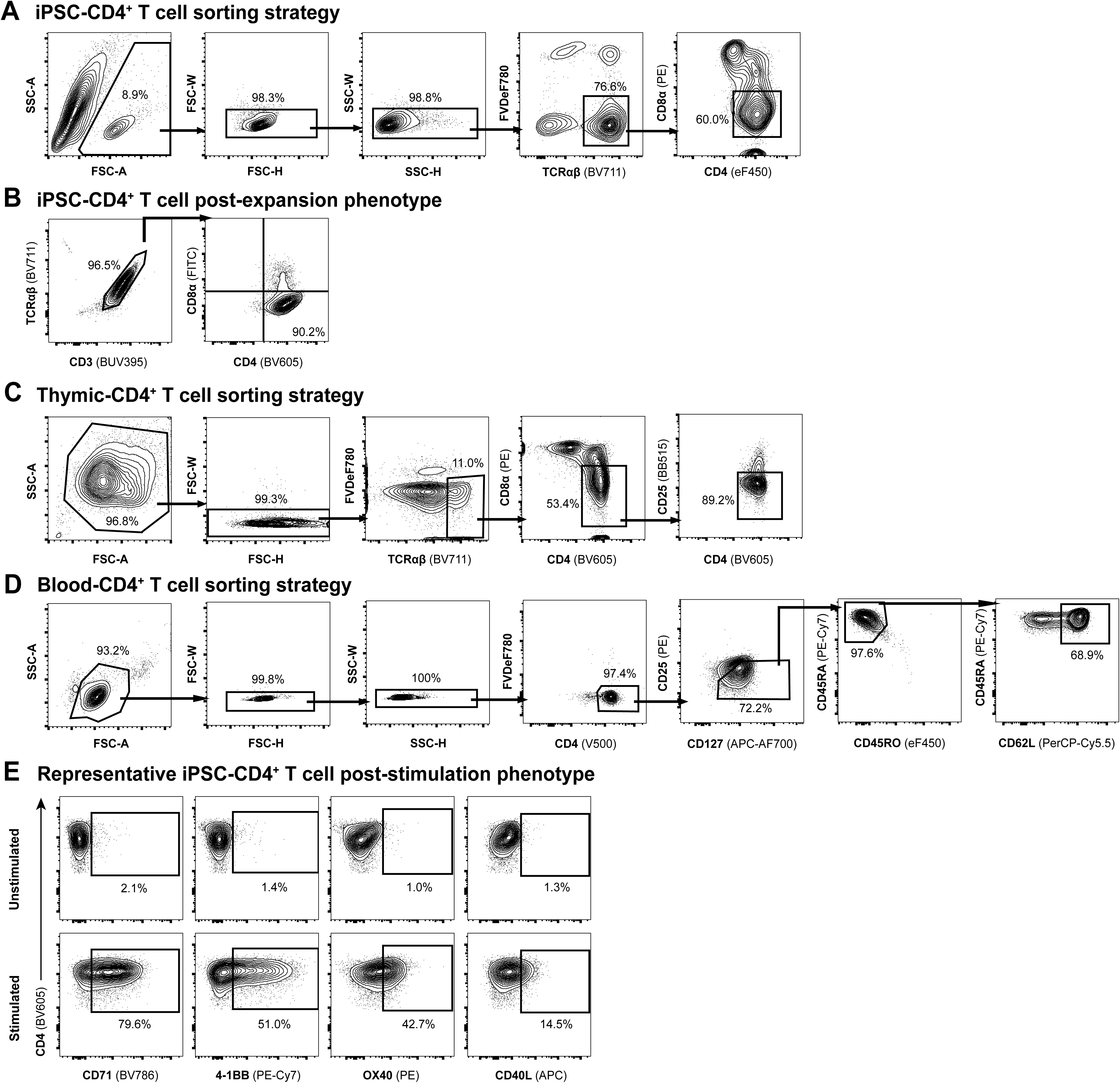
Isolation and expansion of CD4^+^ T cells, related to Figure 3. **(A)** Sorting strategy for isolating iPSC-CD4^+^ T cells. **(B)** iPSC-CD4^+^ T cells were expanded for 14 days, rested overnight in 10 IU/mL IL-2, then measured for percentage of CD4^+^CD8a^-^ within CD3^+^TCRab^+^. Representative flow plots for iPSC-CD4^+^ T cells from Figure 3D on day 15. **(C)** Sorting strategy for isolating thymic-CD4^+^ T cells from human postnatal thymus. **(D)** Sorting strategy for isolating naïve blood-CD4^+^ T cells from leukapheresis products donated by healthy donors. **(E)** Representative flow plots for data from Figure 3E showing activation marker expression in iPSC-CD4^+^ T cells stimulated with anti-CD2/3/28 for 48 hours vs unstimulated.

**Figure S5.**
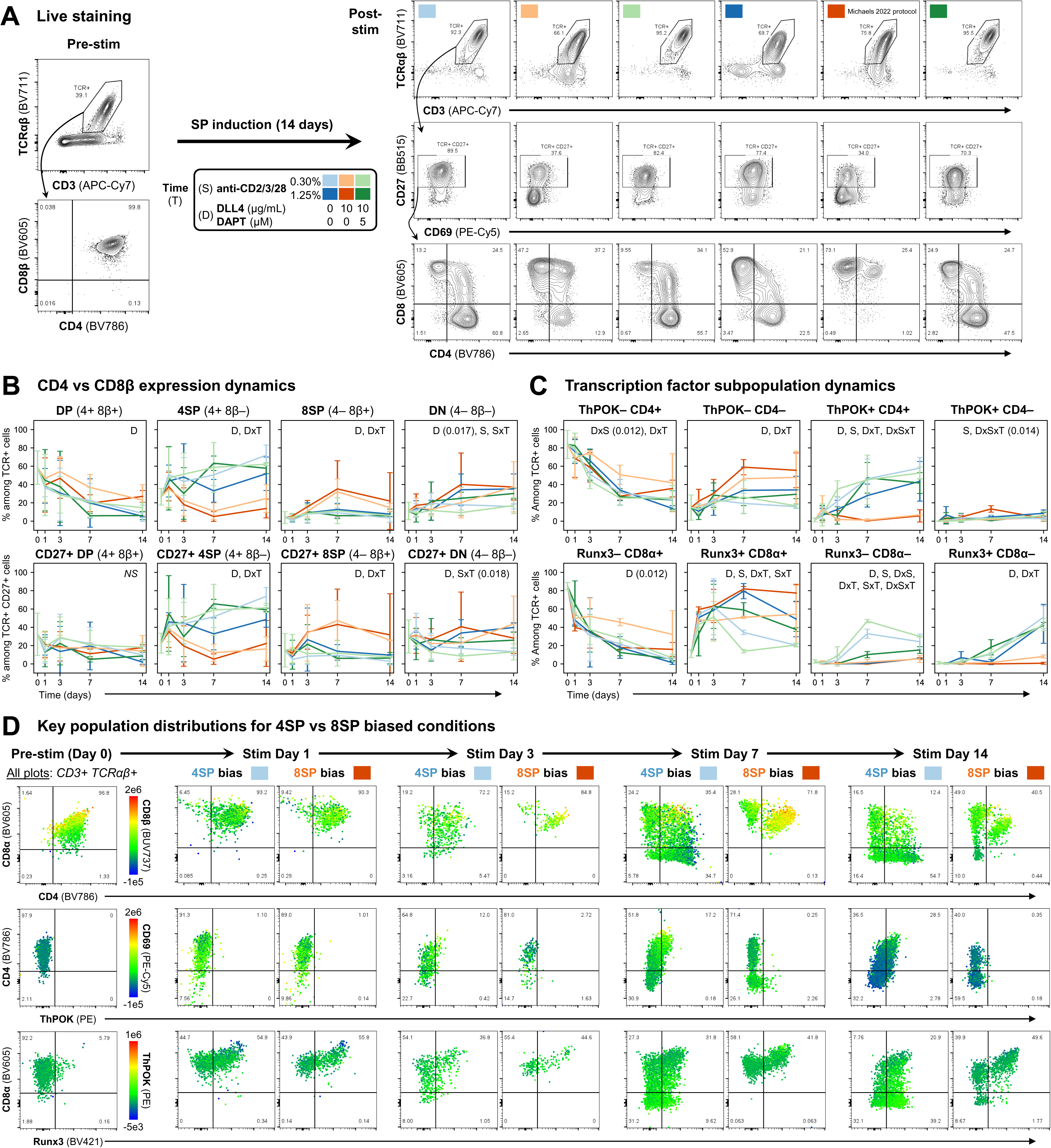
Additional data for TCR stimulation + intracellular staining timecourse, related toFigure 4. **(A)** Representative cells pre-and post-stimulation (day 14) in each different condition. Cells here were stained only with surface markers and not fixed, to provide a live reference for cells fixed and stained for the TFs, after which cellularity drops significantly. **(B-C)** Dynamics of CD4 vs CD8β subpopulations (B) and ThPOK vs CD4 and Runx3 vs CD8α (C). Errorbars: mean ± standard deviation; n=4 independent differentiations (n=2 at day S14); media: StemSpan Mat + JAC Ultra; Inset indicators D, S, and T refer to significance of DLL4/DAPT variants (D), anti-CD2/3/28 levels (S), and interactions (x) with each other and time (T) using three-way ANOVA with a significance threshold of p < 0.01; p-values close to but above the significance threshold are shown. **(D)** Representative distributions of marker expression over time in the main CD4 and CD8 lineage biasing conditions.

**Figure S6.**
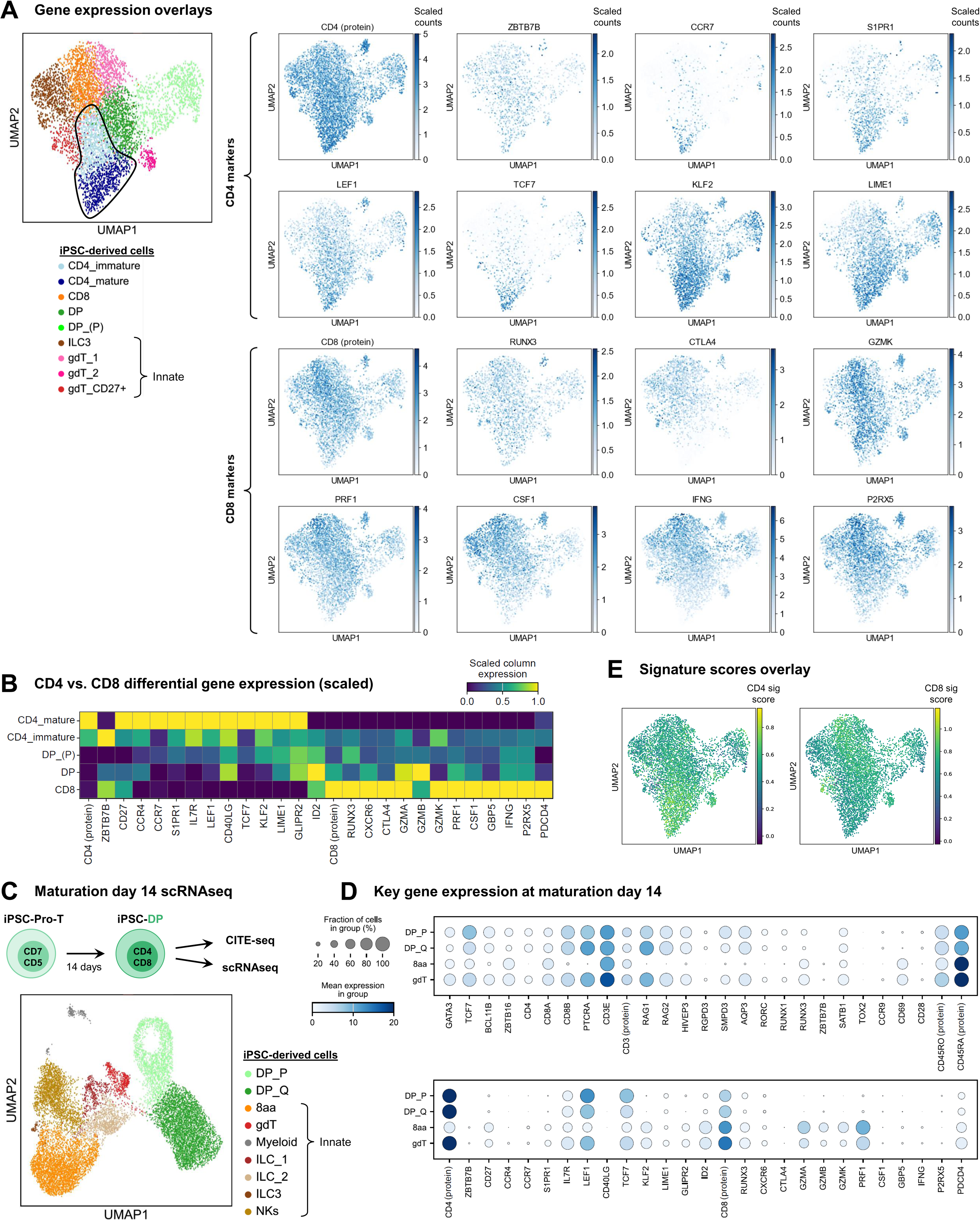
Differential gene expression in iPSC-CD4^+^ vs CD8^+^ T cells, related to Figure 5. **(A)** Overlay of gene expression levels on UMAP space. DP_(P): proliferating DPs; gdT: γδ T cells; ILC: innate lymphoid cells. **(B)** Column-scaled gene expression levels for key genes corresponding to Figure 5C. **(C)** iPSC-derived cells after 2 weeks of maturation and prior to SP induction (day M14, media: PSC2) were collected and sequenced by scRNAseq + CITE-seq. UMAPs and RNA leiden-based clusters identified DP cells (P: proliferating, Q: quiescent), CD8αα innate T cells, γδ T cells, ILC1-, 2-, and 3-like cells, and NK cells. **(D)** Confirmation that unsignaled iPSC-DP cells (i.e. cells pre-TCR stimulation) express expected DP cell marker genes (e.g. *RAG1/2*, *SMPD3*, *AQP3*, *RORC*, and *SATB1*), lack expression of positive selection markers (e.g. *CD69*, *CCR9*, *TOX2*), and express low levels of lineage-specific genes (e.g. *ThPOK*, *RUNX3*, *CD40L*, *GZMA/B/K*), and maturation markers (e.g. *CD27*, *CCR7*, and most others seen in cells post-SP induction and expansion in Figure 5C & **S6B**). **(E)** Overlay of CD4 and CD8 signature scores on UMAP space.

**Figure S7.**
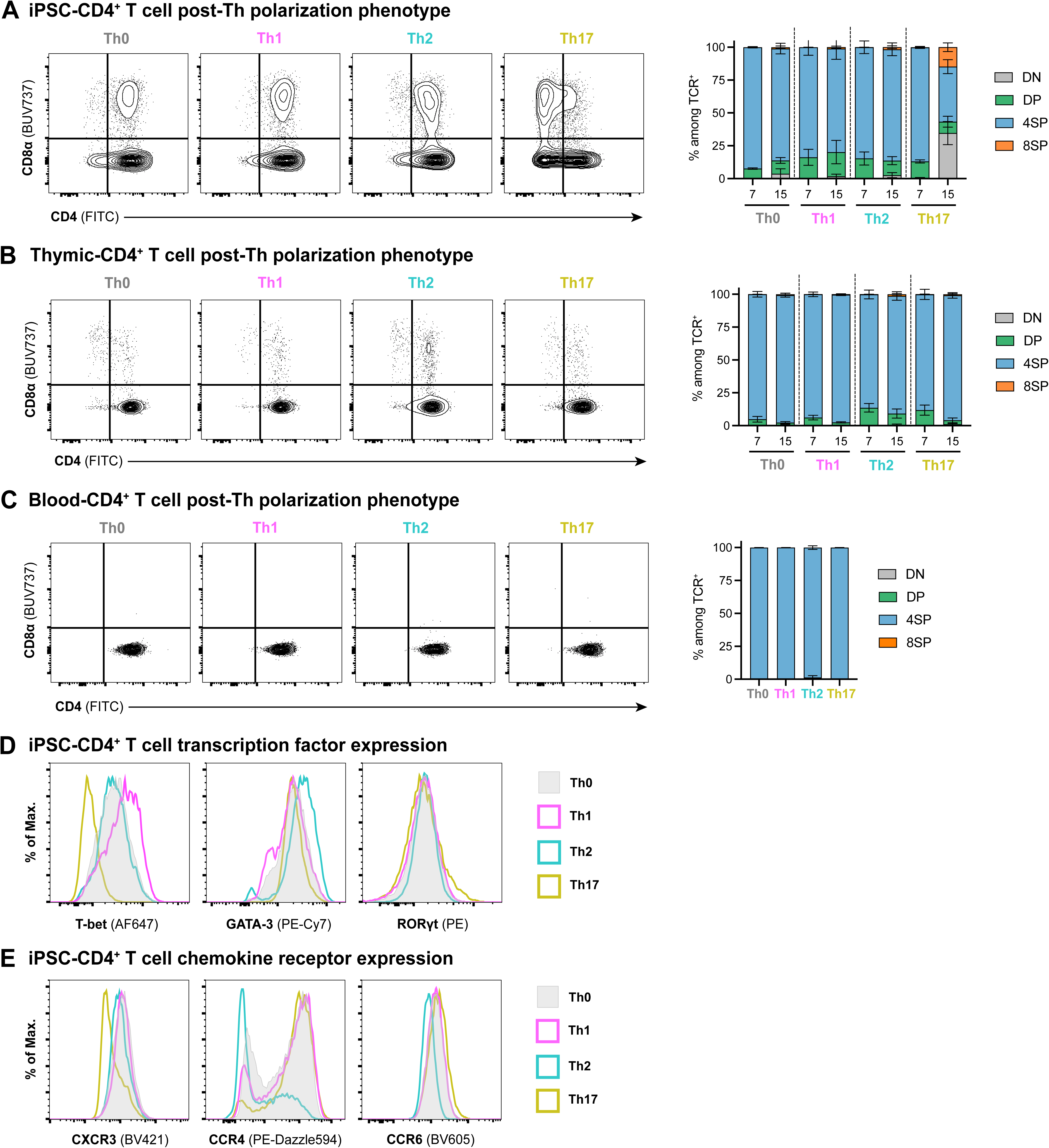
Th stability and cytokine secretion, related to Figure 7. iPSC-, thymic-, and blood-CD4^+^ T cells were polarized in Th1/2/17 conditions for 7-14 days, rested overnight in 10 IU/mL IL-2, and then measured for relevant markers. **(A-C)** Representative plots showing CD4 vs CD8a expression (left) and DN/DP/CD4SP/CD8SP proportion (right) of live iPSC (A), thymic (B), and blood (C) cells on day 7 or 15 (iPSC and thymic) or day 8 (blood). **(D-E)** Representative histograms of rested polarized iPSC-CD4^+^ T cells from Figure 7B-C showing the gMFI of transcription factors (D) and chemokine receptors (E). Data is shown with mean ± standard deviation; n=4, with each symbol representing an individual T cell differentiation or donor (for thymic-and blood-derived cells), collected in 3-4 individual experiments; statistical significance was determined using ordinary one-way ANOVA with p values shown.

## Notes

https://data.mendeley.com/preview/k3xtwnzpyn?a=3209888b-4a90-49f2-b452-77b7d2b4834b

